# Alignment of single-cell RNA-seq samples without over-correction using kernel density matching

**DOI:** 10.1101/2020.01.05.895136

**Authors:** Mengjie Chen, Qi Zhan, Zepeng Mu, Lili Wang, Zhaohui Zheng, Jinlin Miao, Ping Zhu, Yang I Li

## Abstract

Single-cell RNA sequencing (scRNA-seq) technology is poised to replace bulk cell RNA sequencing for most biological and medical applications as it allows users to measure gene expression levels in a cell-type-specific manner. However, data produced by scRNA-seq often exhibit batch effects that can be specific to a cell-type, to a sample, or to an experiment, which prevent integration or comparisons across multiple experiments. Here, we present `Dmatch`, a method that leverages an external expression atlas of human primary cells and kernel density matching to align multiple scRNA-seq experiments for downstream biological analysis. `Dmatch` facilitates alignment of scRNA-seq datasets with cell-types that may overlap only partially, and thus allows integration of multiple distinct scRNA-seq experiments to extract biological insights. In simulation, `Dmatch` compares favorably to other alignment methods, both in terms of reducing sample-specific clustering, and in terms of avoiding over-correction. When applied to scRNA-seq data collected from clinical samples in a healthy individual and five autoimmune disease patients, `Dmatch` enabled cell-type-specific differential gene expression comparisons across biopsy sites and disease conditions, and uncovered a shared population of pro-inflammatory monocytes across biopsy sites in RA patients. We further show that `Dmatch` increases the number of eQTLs mapped from population scRNA-seq data. `Dmatch` is fast, scalable, and improves the utility of scRNA-seq for several important applications. `Dmatch` is freely available online (https://qzhan321.github.io/dmatch/).

## Introduction

Single cell RNA-seq technology is transforming the study of cellular heterogeneity ^1, 2^, differentiation ^3, 4^, and cellular response to stress and stimulation ^5,6,7^. Gene expression levels in tens of thousands of single cells are now routinely measured in a single scRNA-seq experiment ^8^ and more scRNA-seq datasets are becoming available each day. However, there remain considerable challenges in scRNA-seq data pre-processing as gene expression measurements from scRNA-seq are much noisier than compared to bulk RNA sequencing ^9, 10^. This makes integration and comparisons across multiple scRNA-seq experiments particularly difficult because each experiment can vary in capture efficiency, PCR efficiency, and dropout rates, and these technical effects can be cell-type-or experiment-specific ^11, 12^.

Without the ability to integrate multiple scRNA-seq experiments, scRNA-seq studies are limited to two general applications: (1) to characterize cell-type heterogeneity in a population of cells from one experiment ^1,2,8^, or (2) to infer cellular trajectory during development or response to stimuli from one sample. While we have learned tremendously about cellular heterogeneity and cellular state transitions from these studies, there are important applications in both basic and clinical science that require integration and comparisons across multiple scRNA-seq experiments ^11, 12^. Important applications that use scRNA-seq data include the identification of differentially expressed genes between two biological conditions, and the mapping of expression quantitative trait loci (eQTLs).

Here, we describe `Dmatch`, a method that enables integration and comparisons across multiple scRNA-seq experiments. `Dmatch` uses an external panel of primary cells ^13^ to identify shared pseudo cell-types across scRNA-seq samples, and then finds a set of common alignment parameters that minimize gene expression level differences between cells that are determined to be the same pseudo cell-types (Fig. 1a). Finally, `Dmatch` applies an affine transformation in dimensionality reduced space to all gene expression measurements to remove batch or nonbiological effects (Fig. 1a). The affine transformation used by `Dmatch` preserves cell-to-cell relationships and overall structure among cells, and, importantly, retains the cell densities of the original datasets. Thus, aligned data produced by `Dmatch` is well suited for downstream analyses and prevents reduction of cellular variation, which can lead to inflated differential gene expression tests, false positives, and false negatives.

**Figure 1:**
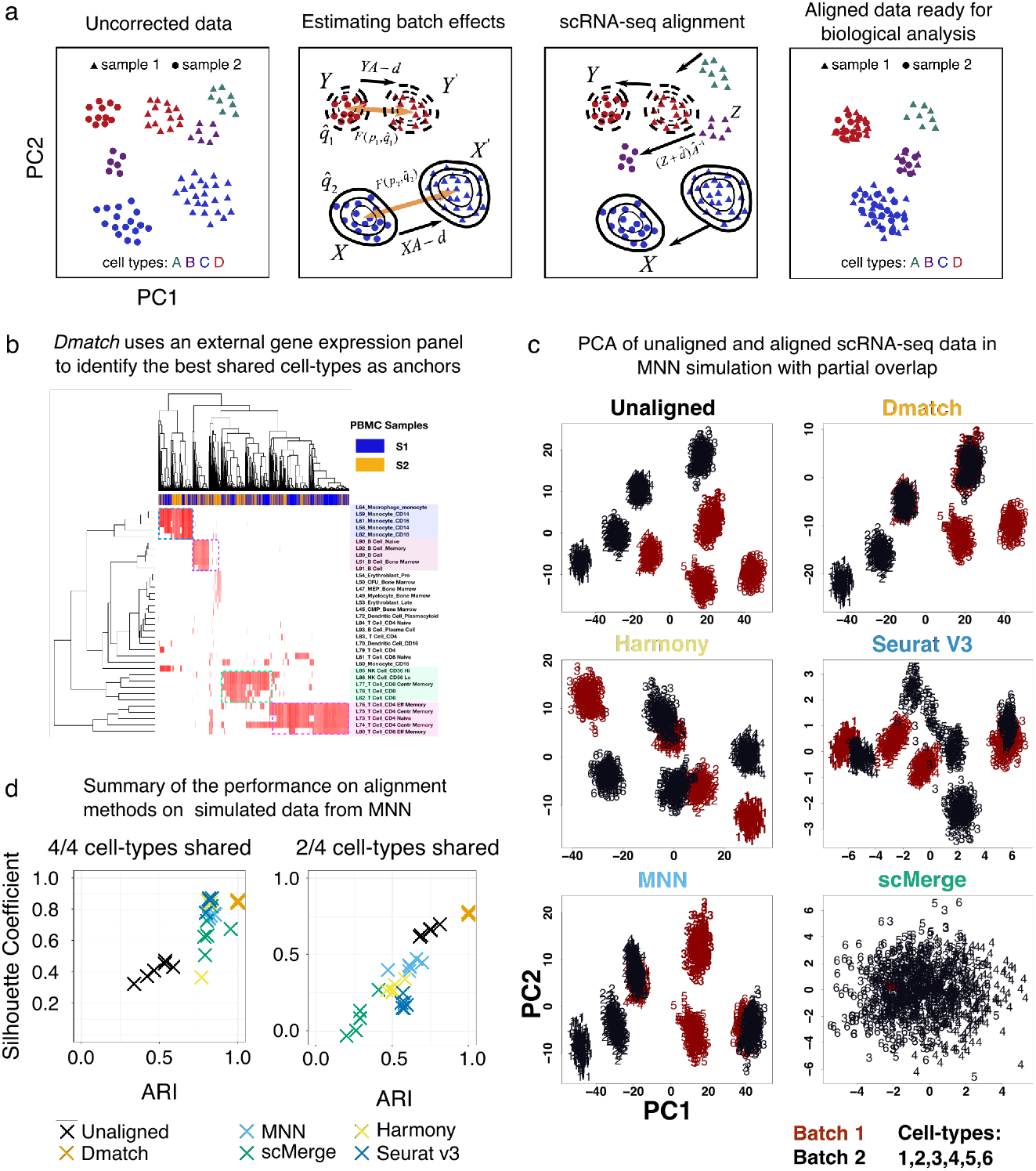
**(a)** Data processing pipeline. First, the uncorrected data is projected onto principal components (PC). Next, an external gene expression panel is used to identify anchor cells to estimate linear batch effects in the form of a rotation and a translation in PC space. Last, the data are corrected by rotating and translating the data points in PC space. The PC loadings are used to recover the aligned data to allow downstream analyses. **(b)** `Dmatch` uses a large reference transcriptomes from the Primary Cell Atlas to identify subpopulations from the observed cells based on the projection. These subpopulations are used as anchors to guide the alignment. We show an example applied on real data, which demonstrates the identification of cell clusters corresponding to monocytes, B cells, and two different subclass of T cells. **(c)** Scatter plot of the first two PCs of unaligned and aligned simulated data using multiple alignment methods. We observed that `Dmatch` was the only method that is was able to accurately align samples with drastic differences in cell-type composition. **(d)** Scatter plot showing two alignment performance metrics, the ARI and the Silhouette coefficient, for unaligned simulated data and simulated data corrected using multiple alignment methods. Simulations were based on the framework proposed previously. We performed two sets of simulations. In the first set, four cell-types were simulated, all of which were shared across the samples to be aligned. In the second set, only two cell-types out of four were shared across the samples.

We showcase `Dmatch` on two major applications including differential gene expression analysis of cell clusters from biopsies from healthy and disease individuals, and cell-type-specific eQTL mapping. `Dmatch` compares similarly or favorably to existing methods on evaluation metrics. Most notably, `Dmatch` is designed to integrate scRNA-seq data without removing biological variation and we show that `Dmatch` is less prone to overcorrection compared to existing methods.

## Results

### Identification of shared cell-types across scRNA-seq experiments as anchors

To identify shared cell-types across a pair of scRNA-seq datasets, `Dmatch` computes the Pearson correlation between gene expression levels of all cells and gene expression quantifications of primary cells from the Primary Cell Atlas ^13^. More specifically, the Primary Cell Atlas is a meta-analysis of publicly available microarray datasets compiled from 95 human primary cells from over 100 separate studies ^13,14^ and was previously used in Reference Component analysis ^15^. Using gene expression measurements from the Cell Atlas, `Dmatch` computes, for each cell in the scRNA-seq experiments to be aligned, a 95-dimensional vector of Pearson correlations – one correlation per primary cell. Because cells with similar Pearson correlation vectors are more likely to correspond to the same biological cell-types, we reasoned that cells from different scRNA-seq experiments with similar correlation vectors likely belong to the same biological clusters. `Dmatch` thus clusters cells based on their Pearson correlation vector, and uses these clusters as anchors to determine alignment parameters.

To improve the consistency of the cell clustering, we implemented various strategies to reduce noise in cell-type assignment and to detect outlier cells in scRNA-seq datasets. For example, based on empirical tests, we found that the consistency of cell-type assignment was generally increased when all Pearson correlations of the 95-dimensional vectors were set to zero except for those between the cell and the top five reference atlas cell-types with the highest Pearson correlation coefficient (the results were qualitatively similar when the top five to ten reference cell-types were used). To minimize the chance that a rare cell-type from the reference panel is chosen as a reference cell-type, we required that all reference cell-types considered must be ranked as a top five highest correlated cell-type with more than 20 cells from the scRNA-seq datasets. Altogether, this procedure results in a sparse Pearson correlation matrix, which is then biclustered (e.g. Fig. 1b, Methods). We found that inducing sparsity in this matrix reduced the noise in clustering, which became immediately visible (see Supplementary Figure 1 for comparison). `Dmatch` also uses two criteria to rank cell clusters and select two or more clusters as anchors. First, `Dmatch` favors clusters with at least 100 cells in each scRNA-seq sample. Second, `Dmatch` performs a Shapiro-Wilk test to rank cell clusters in terms of their likelihood to be drawn from a normal distribution in PC space, an assumption that is required in our optimization (described below). The normality assumption also helps detect cell clusters that visibly consist of two or more distinct cell sub-clusters or clusters with a large proportion of outlier cells. Clusters that fail the Shapiro-Wilk test are removed from the pool of possible anchors.

### Alignment of scRNA-seq data using anchor cell-types

`Dmatch` uses two or more shared cell-types as anchors to estimate alignment parameters to reduce cell-type-specific batch effects. Specifically, `Dmatch` first estimates the probability density distribution of anchor cells in each sample separately using Gaussian distributions to model each probability density. Then, `Dmatch` uses gradient descent to find the best linear transformation (translation *d* and rotation *A*) to minimize the Kullback-Leibler Divergence (KL divergence) between cells from anchor cell-types in one “source” scRNA-seq dataset and the matched cells from another “target” dataset (Fig. 1a, Methods). Once the parameters that minimize the KL divergence are obtained, the linear transformation is applied to all cells, including non-anchor cell-types, from the “source” scRNA-seq dataset. The transformed values from the “source” dataset, combined with the original values from the “target” dataset, form a pairwise alignment between the two scRNA-seq samples. This alignment process can be repeated iteratively to align additional scRNA-seq samples to the “target” dataset.

Although our approach is applicable when only a single anchor cell-type is shared across two scRNA-seq experiments, we recommend using at least two anchors to reduce the potential for over-correction and introducing cell-type-specific batch effects. Indeed, batch effects have been observed to depend on mRNA expression level, length, and nucleotide composition ^16, 11^. Thus, batch effects can also be cell-type-specific because the expression levels of genes and the amount of total mRNA molecules vary across cell-types. By estimating alignment parameters using two or more anchors drastically reduces over-correcting cell-type-specific batch effects, which prevents artificial signals to be introduced in the aligned data. To evaluate the consistency of our alignment when different cell clusters are selected as anchors, we computed the mean squared error (MSE) between datasets produced using different anchors (Methods). We found that the MSE between datasets aligned using different anchors were significantly smaller (average 0.59) than compared to the MSE between unaligned data and aligned data (average 9.74), suggesting that choosing different anchors result in similar alignments.

### Evaluation of alignment methods on simulated data

We evaluated the performance of `Dmatch` and four recently proposed alignment methods, including Seurat V3 ^17^, MNN (scran 1.9.39) ^11^, scMerge ^18^(0.1.9.1), and Harmony ^19^ (0.0.0.9000). We first applied all methods on simulated scRNA-seq data. Our simulated samples were assumed to follow a four-component Gaussian mixture model in two dimensions to represent gene expression profiles of four cell-types in a low dimensional biological subspace (Methods). Two samples each with 1,000 cells were drawn with mixing coefficient (0.25, 0.25, 0.25, 0.25) as the first and second batch, respectively. Furthermore, two scenarios with different levels of difficulties were considered. In the simple case, all four cell-types are shared. In the difficult case, batches are partially overlapping with only two cell-types shared. We used data generating models from the MNN software ^11^ to generate simulated datasets. Specifically, artificial batch effects were added to one dataset by adding a Gaussian random vector to the expression profiles of all cells in that dataset. This dataset was then projected onto a 100 dimensional-space using a matrix of values that corresponds to a subset of the principal component loadings obtained from real data. The resulting vectors simulate high-dimensional gene expression data.

In the simple case, we observed that all methods performed similarly well (Supplementary Figure 2). However, only `Dmatch` correctly merged cell-types that were shared across batches in the difficult case (Fig. 1c), while the other methods tended to either over-correct or under-correct batch effects. Our simulation thus shows that `Dmatch` can robustly correct batch effects that induce systematic linear effects on gene expression levels for partially overlapping samples, even when the batch effects are generated according to the MNN model.

We also evaluated the performance of all methods measured using two popular metrics: the silhouette coefficient ^20^ and Adjusted Rand Index (ARI) ^21^ (Methods). The Silhouette coefficient aims to evaluate how well mixed the cells which belong to the same cell-type across batches versus how well separated the cells which belong to different cell-types. ARI evaluates the consistency between the true cell labels versus the assigned cell labels after batch effects correction. In the easy case (Fig. 1d, 4/4 cell-types shared), we found that `Dmatch` achieved an average SC of 0.85. Seurat V3 and Harmony achieved an average SC of 0.85 and 0.77, respectively, while MNN and scMerge performed the worst. In terms of ARI, `Dmatch` assigned labels that matched the true labels in all simulations and reached a score of 1. By contrast, the other methods obtained an average ARI between 0.7 and 0.9. In the difficult case (Fig. 1d, 2/4 cell-types shared), `Dmatch` achieved an average SC of 0.77, whereas the other methods scored below 0.5. Similarly, `Dmatch` achieved a ARI score of 1, whereas other methods scored below 0.5.

### Alignment of clinical scRNA-seq data from patient biopsies

To evaluate the five alignment methods on real data, we chose to focus on immune cells as they have been characterized extensively, and many cell-types have well-documented markers. To obtain a realistic dataset that uses the same scRNA-seq technology to measure gene expression levels at different biopsy sites in multiple individuals, we collected scRNA-seq data using 10X genomics (Methods) from PBMC (average 4,444 cells) and bone marrow (average 1,013 cells) from six individuals (12 total samples), including three rheumatoid arthritis (RA) patients, one ankylosing spondylitis (AS) patient, one systemic lupus erythematosus (SLE) patient, and one healthy individual (HC) (Supplementary Table 1). In two of the three RA patients, we further collected scRNA-seq data from synovial fluid (2,961 and 2,254 cells) at the active site of inflammation. Although these samples were collected and processed on the same day, they were processed on different experimental runs and were not multiplexed nor pooled together. Thus, our data collection reflects a realistic collection procedure for clinical samples that we predict will be widespread.

We first aligned the six PBMC samples we collected to survey the general landscape of immune cells in healthy and disease peripheral blood. Using UMAP to visualize the unaligned data, we observed subtle, but clear, separation between clusters of cells from different individual samples, indicative of the presence of batch effects (Figure 2a). As expected, all alignment methods produced a noticeable reduction in the separation of cells from different samples. The mixing of sample cells after alignment using Harmony and in particular scMerge was particularly homogeneous (Figure 2a).

**Figure 2:**
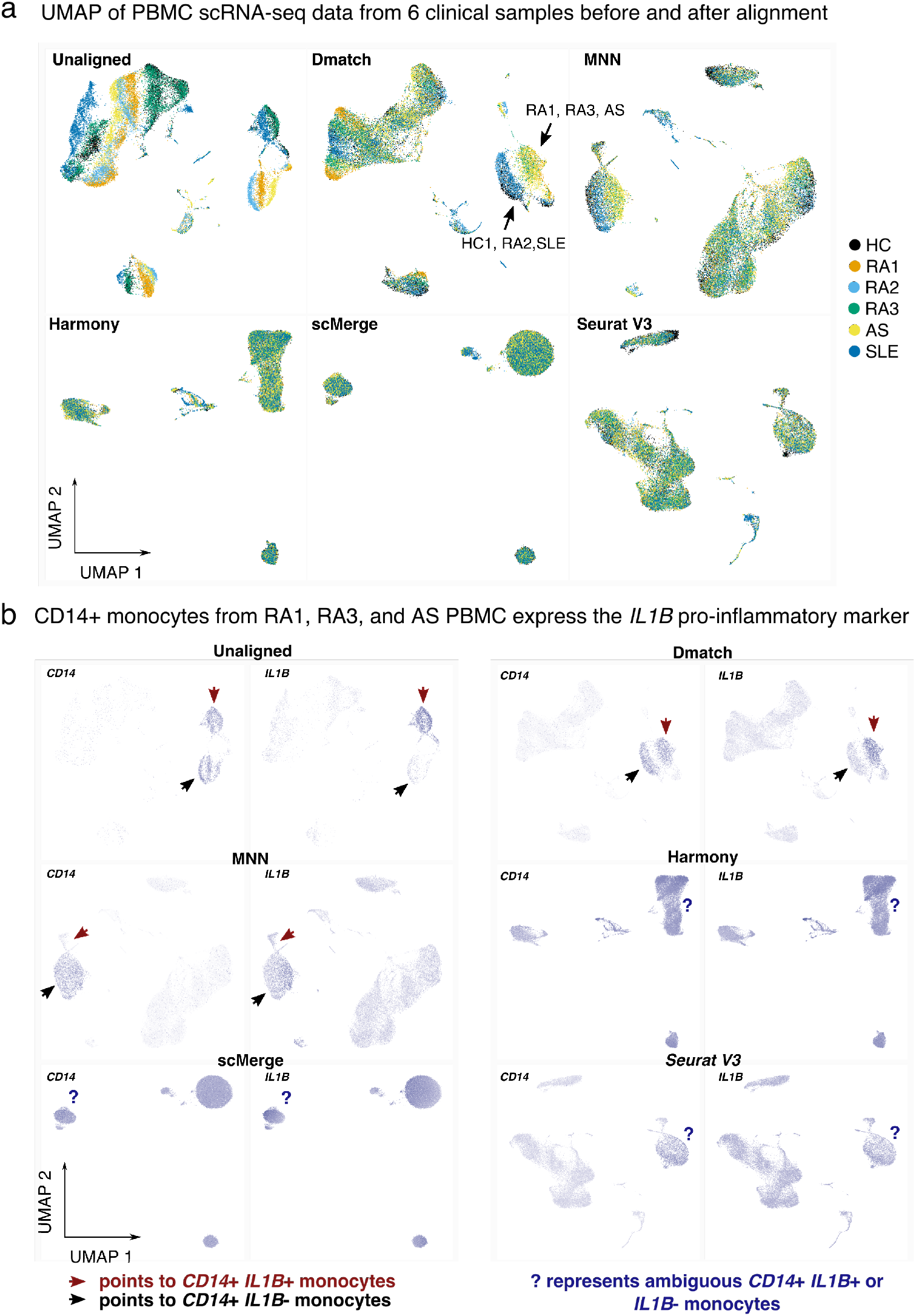
Application of alignment tools on PBMC clinical samples. **(a)** UMAP dimensionality reduction of PBMC scRNA-seq data from 6 clinical samples before and after alignment comparing `Dmatch`, MNN, Harmony, scMerge, and Seurat V3. UMAP plots suggest possible over-correction from scMerge. **(b)** We observed that *CD14* monocytes express *IL1B* in a subset of individuals with autoimmune disease (RA1, RA3, AS). This signal was preserved after alignment using `Dmatch` and to some extent MNN, while it disappeared using Harmony, Seurat V3, and scMerge. Shades of purple represent marker expression in each cell relative to other cells in the same UMAP.

To characterize more systematically the accuracy of the alignment, we sought to determine whether the same cell-types from different samples were aligned together while distinct cell-types remained separated. To this end, we applied a standard pipeline to identify cell clusters using Seurat on each sample separately (Method). This resulted in 8–15 clusters for the 6 samples, wherein each cluster expressed informative markers that allowed us to assign a likely immune cell-type (e.g. *CD4*+ T cell, or *CD14*+ monocyte, see Methods). Visualizing these markers on the UMAP revealed that, indeed, the alignment procedures were able to align cells expressing the same markers together (Figure 2a and Supplementary Figure 3). However, in addition to aligning similar cell-types together, scMerge also aligned distinct cell-types together, resulting in three major homogeneous cell clusters representing B cells, monocytes, and a large cluster of T cells (Supplementary Figure 3).

When the samples were analyzed separately using the standard Seurat pipeline (Methods), we observed that *CD14*+ monocytes from RA1, RA3, and AS (but not RA2, SLE, or HC) expressed the pro-inflammatory marker *IL1B*, indicating that monocytes in these individuals exhibit a pro-inflammatory state. *IL1B* positive *CD14*+ monocytes have been observed in RA patients ^22^, but the expression of *IL1B* in monocytes from the ankylosing spondylitis patient support reports that *IL1B* may be involved in the pathogenesis of a wide number of autoimmune diseases ^23, 24^.

We also observed a cluster in SLE PBMC consisting of cells that expressed high levels of kidney-expressed genes (e.g. *ECHS1*, *MIOX*, *FXYD2*, and *ALDOB*). The presence of this “kidney” cell cluster is consistent with circulating kidney cells in the peripheral blood of the SLE patient as the patient exhibits kidney inflammation in the form of lupus nephritis.

Because both pro-inflammatory monocyte clusters and the “kidney” cell cluster were identified in the patient samples when analyzed individually, we reasoned that their presence could not be explained by technical effects. However, investigating data aligned using Seurat v3, Harmony, and scMerge revealed that the clusters representing pro-inflammatory monocytes disappeared subsequent to alignment (Figure 2b) and the cluster corresponding to SLE-specific circulating kidney cells disappeared after Seurat V3, Harmony, and to some extent, scMerge alignment (Supplementary Figure 4). By contrast, both clusters are visibly separated in the UMAP representation of data aligned using `Dmatch` and, to some extent, MNN (Figure 2b and Supplementary Figure 4).

### Over-correction of batch effect masks biological signal

To better understand the extent to which data aligned using different methods vary, we performed a series of differential gene expression analyses between cell-type clusters that were determined using Seurat on each sample separately, henceforth referred to as Seurat clusters (see Methods). We thus set to identify differences in the list of genes identified as differentially expressed across Seurat clusters when different methods were used to align the data. We reasoned that if the lists of differentially expressed genes (DEG) across Seurat clusters are the same for all methods, then the alignments must be very similar across methods. However, if the lists of DEG differ dramatically, then the alignments must be highly variable.

We began by evaluating the differences between unaligned data and data aligned using `Dmatch`. For each pair of Seurat clusters, we counted the number of DEG that were identified using limma-trend ^25^ (see Methods) using unaligned data as input but not using data aligned from `Dmatch` as input, and vice versa. We first identified DEGs across pairs of Seurat clusters from different samples but that were determined to represent the same cell-type from our manual annotation (Methods). In the ideal scenario, the number of DEG between clusters representing the same cell-type should be reduced when using aligned data compared to unaligned data, because DEG identified when comparing the same cell-type across different samples are generally expected to be due to batch effects. Indeed, when we compared DEG obtained from unaligned and `Dmatch`-aligned data, we found that the number of DEG between same cell-types in different samples were drastically reduced (generally 5–25 DEGs from unaligned data versus 0-3 DEGs from `Dmatch`-aligned data) (Figure 3a, X’s). The reduction in the number of DEGs therefore suggests that `Dmatch` was able to efficiently remove the effects of batch.

**Figure 3:**
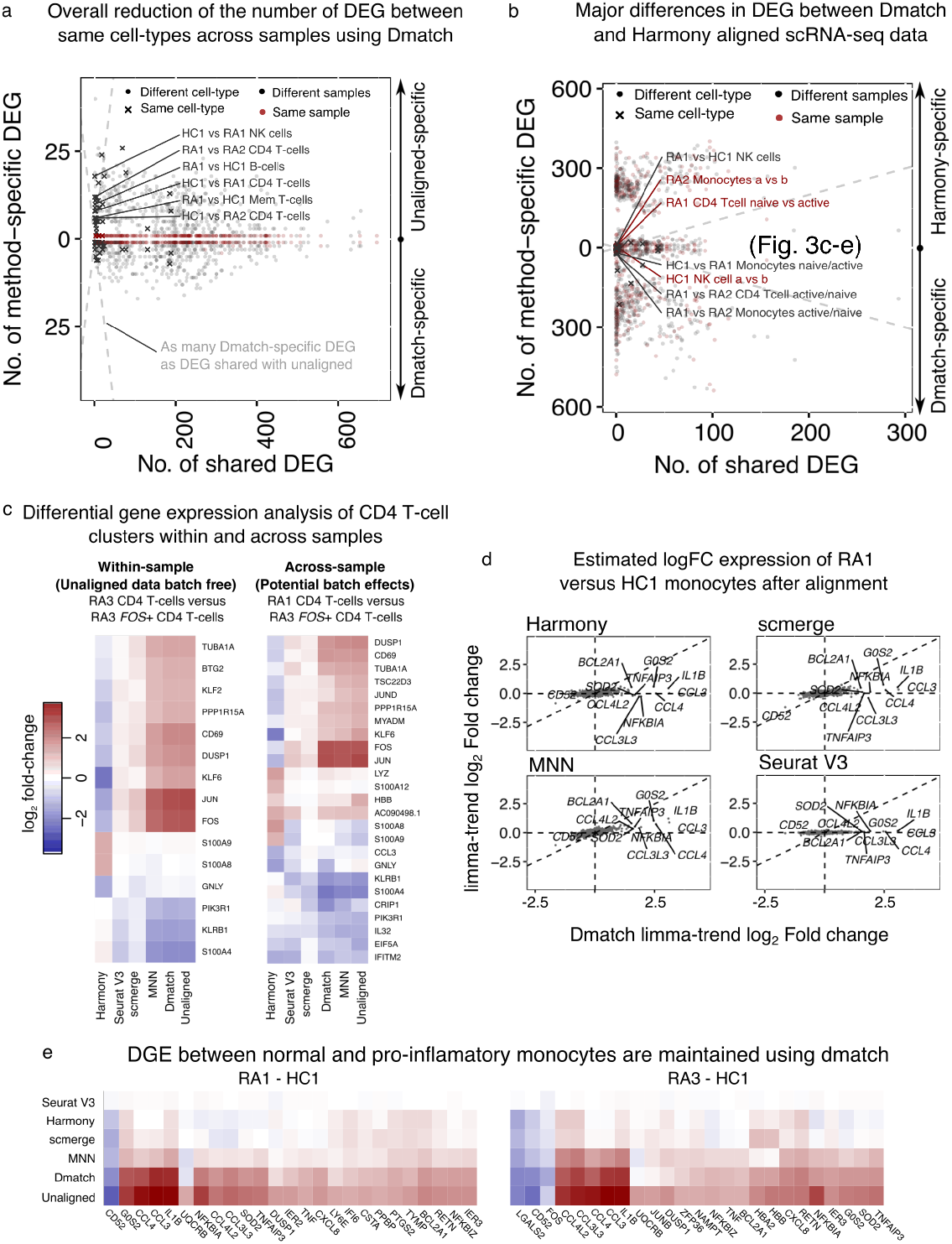
Over-correction in alignment tools. **(a)** Scatter plot showing the number of differentially expressed genes (DEG) that are identified using unaligned data (positive y-axis) or using data aligned using `Dmatch` (negative y-axis), versus the number of DEG that are identified in both datasets (x-axis). Each point represents a comparison between cell-type clusters from the same or different cell-type, or from the same or different sample. Successful removal of batch effect is supported by smaller numbers of DEGs resulting from `Dmatch`-aligned data in same cell-type comparisons, compared to unaligned data. As expected, DEG inferred from within-sample cluster comparisons are identical between unaligned and `Dmatch`-aligned data. **(b)** Scatter plot similar to (a) shows a large difference between `Dmatch` and Harmony-aligned data. **(c)** Heatmaps of estimated DEG fold changes between naive *CD4* T cells versus activated *CD4* T cells within the same sample (RA3, left heatmap), and across two samples (RA1 vs RA3, right heatmap). `Dmatch` and MNN estimates of DEG fold changes within samples are consistent with unaligned data, as we should expect. However, the estimates from scMerge and Harmony are shrunken and inconsistent, respectively, suggesting over-correction. Across-sample comparison between the two cell-types show increase in *JUN*, *FOS*, and *CD69* expression in activated CD4+ T cells, consistent with within-sample comparisons for `Dmatch` and MNN. The signal is reduced or reverse for scMerge and Harmony, respectively. **(d)** Scatter plot of DEG log fold change comparison between CD14 monocytes from healthy individual HC and RA patient (RA1). Fold change estimates are reduced or zero for inflammatory markers (*IL1B*, *CCL3*, *CCL4*) using data obtained from MNN, Harmony, and scMerge. **(e)** Heatmaps of estimated DEG fold change comparison between RA1-HC1 and RA3-HC1 are consistent, and support the presence of over-correction in data aligned using Harmony, scMerge, and MNN, masking real biological signal.

We also considered comparisons across pairs of Seurat clusters determined to be different cell-types, but from the same sample (Figure 3a and 3b, red points). Ideally in this case, DEGs identified across two clusters from the same sample after alignment should be largely the same as those identified before alignment because much of the confounding effects are shared among measurements in a single experiment. In fact, within-sample DEGs identified from unaligned data that are not identified from aligned data suggest overcorrection and a reduction of true biological variation. Reassuringly, we found that DEGs from within sample comparisons were identical between unaligned data and data aligned using `Dmatch`. Altogether, these results suggest that `Dmatch` is able to preserve within sample biological variation while correcting across-sample batch effects.

We next compared DEGs identified from unaligned data to DEGs identified from Harmony, MNN, scMerge, and Seurat V3. We found that while all methods showed reduction of DEGs identified across Seurat clusters determined to the same cell-types, there were also significant differences in the DEGs found between Seurat clusters within a sample (Supplementary Figure 5). These observations raise the possibility that existing alignment methods are prone to overcorrection.

We next compared DEG results between `Dmatch` and other methods. We found drastic differences between the DEGs inferred from aligned data (e.g. between `Dmatch` and Harmony, Figure 3b, for other comparisons see Supplementary Figure 6). For example, many comparisons shared no DEGs while hundreds of genes were identified as differentially expressed in either `Dmatch`-aligned data only or Harmony-aligned data only. Although the number of DEGs in comparisons between the same cell-types was smaller using Harmony and some of the other methods, the large number of method-specific DEGs when comparing clusters within samples suggests overcorrection.

To further investigate possible overcorrection, we focused on identifying cell clusters that show biologically-relevant or known differences. We found that samples from RA1 and RA3 both harbored two clusters of cells that expressed classical markers of CD4+ T cells (e.g. *CD3D*, *IL7R*, *IL32*), with one cluster showing high expression levels of activation markers including *FOS* and *JUN* (Supplementary Figure 7). Because both clusters were found in RA1 and RA3 patients, they likely represent real biological variation between naive and activated CD4+ T cells circulating in patient blood. To verify this, we identified DEG between the naive and activated CD4+ T cell clusters. As expected, within the top 15 most significantly DEG, we observed over-expression of activation makers such as *CD69*, *JUN*, *FOS*, *DUSP1* in activated T cells when using unaligned data, `Dmatch`-aligned, and MNN-aligned data (Figure 3c). However, we found that the estimated fold changes were reduced or even opposite when using Seurat V3, scMerge or Harmony aligned data (Figure 3c). We also observed that comparing naive CD4 T cells from RA1 to activated CD4 T cells from RA3 resulted in a highly consistent list of DEG to within sample comparison, further suggesting that cross-sample comparison is likely accurate for MNN and `Dmatch`, and inaccurate for Seurat V3, scMerge and Harmony.

We also found that the list of DEG between the CD14+ monocytes Seurat cluster from RA1 and the CD14+ monocytes Seurat cluster from HC were different between `Dmatch`-aligned data and data aligned using other methods. As described earlier, we found that the pro-inflammatory marker *IL1B* was over-expressed in RA1 CD14+ monocytes compared to monocytes from the healthy individual (HC). We also observed over-expression of additional consistent markers such as *CCL3*, *CCL4*, *NFKBIA* (Figure 3d,e). By contrast, starting from data aligned using MNN, scMerge, Seurat V3, and Harmony, the differences in gene expression levels between RA1 and HC monocytes were drastically reduced in effect sizes (MNN) or disappeared completely (Harmony, Seurat V3, and scMerge) (Figure 3d,e). *IL1B* and cytokines (e.g. *CCL3* and *CCL4*) have long been recognized to be over-expressed in samples from rheumatoid arthritis patients ^26, 23^. Thus, our observations suggest that heterogeneity between the two monocyte clusters are not a result of uncorrected batch effects. Instead, the lack of DEG detected across the two monocyte clusters using Harmony, scMerge, SeuratV3 – and to some extent MNN – indicates that the two biologically variable clusters are aligned together erroneously.

Altogether, these results suggest that `Dmatch` allows correction of batch effects from scRNA-seq data, while avoiding over-correction. In contrast, while Harmony, scMerge, Seurat V3, and MNN are all able to correct batch effects by homogenizing scRNA-seq data from different experiments, they do so at the cost of removing real biological signals.

### Variation in gene expression levels in peripheral and tissue-resident immune cells

We next evaluated scRNA-seq alignment across biopsy sites. To this end, we used all methods to align scRNA-seq data from PBMC and bone marrow (BMMC) from all individuals (12 total samples) and again evaluated the number of DEGs across Seurat clusters. We then conducted a hierarchical clustering on all samples based on the number of DEG as the distance metric. We found that compared to the clustering from unaligned data, the clustering using aligned data from all methods resulted in more clearly defined clusters (Figure 4a, Supplementary Figure 8). As expected, the samples clustered according to inferred cell-type, rather than sample provenance, and samples within the same cell-type clusters showed fewer DEG within cluster than compared to unaligned data, suggesting a reduction of batch and technical effects.

**Figure 4:**
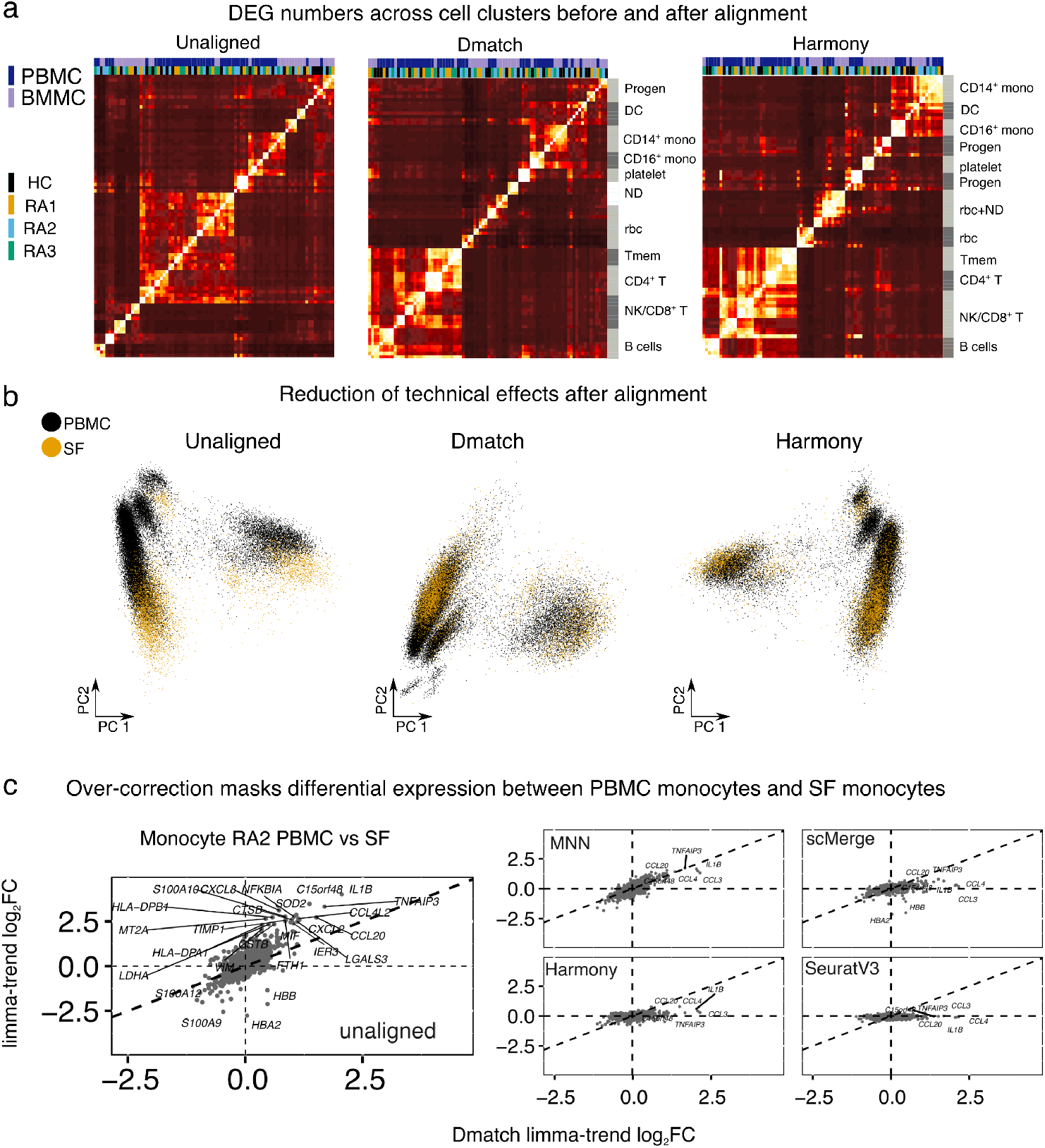
Cross-tissue alignments **(a)** Heatmaps showing the number of DEGs identified (Methods) across Seurat clusters in PBMC and BMMC samples using unaligned data, `Dmatch`-aligned data, and Harmony aligned data (others in Supplementary Figure 8). Improvement of clustering by cell-type using aligned data indicates removal of batch effects from unaligned data. ND: Not Determined. **(b)** PCA of scRNA-seq samples from PBMC and SF before alignment, and after alignment using `Dmatch`, and Harmony. While unaligned data show likely batch effects specific to biopsy site, this effect is reduced after alignment. **(c)** Scatter plots of estimated gene expression log fold changes between *CD14*+ monocytes from RA2 PBMC and RA2 synovial fluid (SF). We found that only `Dmatch`- and MNN-aligned data led to the discovery that several pro-inflammatory genes were over-expressed in RA2 SF monocytes.

The study of differences between peripheral immune cells and resident immune cells in the pathological sites requires integration of peripheral samples and samples from the pathological sites. Because cell-type composition can differ across these sites, we predicted that `Dmatch` would outperform other methods as, based on our simulations, existing methods struggled to align samples in the presence of cell-types that were not shared or high variability in cell-type proportions. To test our prediction, we aligned our scRNA-seq data from synovial fluid (SF) (from RA1 and RA2) to scRNA-seq data from PBMC (all individuals). PC analysis on uncorrected data suggests that SF and PBMC scRNA-seq showed clustering within biopsy site, indicative of the presence of batch effects (Figure 4b). Alignment of the data using `Dmatch` showed clear improvement in mixing of cells from SF and PBMC samples (Figure 4b). Perhaps unexpectedly, we observed that alignment using other tools such as Harmony also showed improved mixing and a clear reduction of batch effects (Figure 4b, Supplementary Figure 9).

To better understand the differences in alignments produced by the five methods, we again compared the list of DEGs across Seurat clusters in PBMC and synovial fluid scRNA-seq dataset without alignment and after alignment using the five methods. We found that using unaligned data generally led to a larger number of DEGs across samples from different biopsy sites (Supplementary Figure 10), as expected from the large batch effects observed from the PCA (Figure 4b).

However, we found that many pairs of cell clusters had DEG that were only identified in `Dmatch` and MNN, but not Harmony, scMerge or SeuratV3. Among the pairs of clusters with DEGs specific to `Dmatch` and MNN are two clusters of monocytes from RA2 PBMC and RA2 SF. Upon examination, we found that while monocytes from RA2 PBMC did not express *IL1B* or other pro-inflammatory markers, *IL1B* and other pro-inflammatory markers were highly over-expressed in RA2 SF monocytes. This is by contrast to RA1 samples, in which pro-inflammatory monocytes were found in both PBMC and SF. Interestingly, `Dmatch`-aligned data revealed that monocytes in RA1 SF expressed higher levels of *CCL3* compared to RA1 PBMC monocytes, suggesting a stronger or more robust activation of monocytes in SF than in PBMC. Our finding suggests that presence of *IL1B*+*CCL3*+*CCL4*+ monocytes may be a potent biomarker of rheumatoid arthritis, even though not always detectable in patient PBMC.

Importantly, we were able to identify *IL1B* over-expression on monocytes only using data aligned by `Dmatch` and MNN – and the extent of the over-expression estimated using MNN alignments was reduced compared to that estimated by `Dmatch`. *IL1B* over-expression and thus this entire population of pro-inflammatory monocytes could not be detected in scRNA-seq data aligned using Harmony, scMerge, and SeuratV3.

Thus, we conclude that while all methods were able to integrate scRNA-seq samples from multiple biopsy sites with modest differences in cell-type compositions, nearly all existing methods also removed true biological variation between cells from the different biopsy sites.

### Alignment improve power of eQTL mapping from population-level single-cell RNA-seq data

PCA is often used to estimate and correct batch effects to increase mapping power in eQTL studies from bulk RNA-seq data ^27^. However, because PCA-based methods often fail to correct batch effects across scRNA-seq samples, we hypothesized that `Dmatch` and other alignment methods can increase eQTL mapping power beyond standard PCA-based batch correction methods. To test this, we re-analyzed population scRNA-seq data from peripheral blood mononuclear cell collected in a previous study ^28^. UMAP of the unaligned data show minimal batch effects (Figure 5a), as has been generally observed from high quality droplet-based scRNA-seq from peripheral blood.

**Figure 5:**
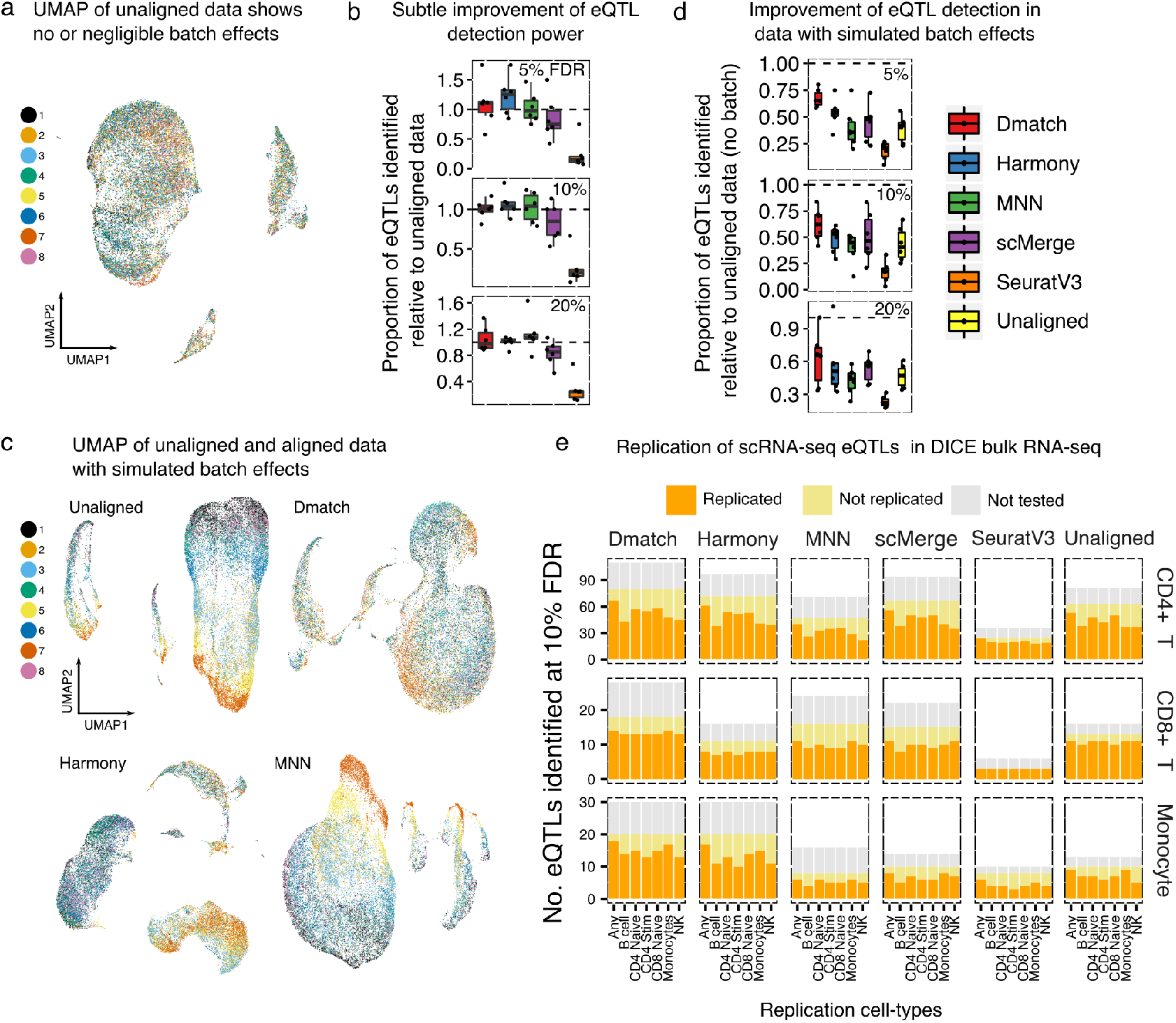
**(a)** UMAP of unaligned PBMC scRNA-seq data from 45 donors ^28^. The cells from the 45 donors were pooled into 8 samples for sequencing. **(b)** Alignment slightly improve mapping power. **(c)** UMAP of unaligned PBMC data with spiked in batch effects (see Methods) **(d)** Alignment improve eQTL mapping power substantially when batch effects are simulated to reflect a realistic and practical scRNA-seq collection strategy. **(e)** Replication of eQTLs identified in a bulk RNA-seq study of immune cell-type eQTLs from the DICE consortium. The eQTLs that were not tested correspond to SNPs whose genotype could not be determined or to genes filtered out due to low expression.

We mapped eQTLs in 8 cell-types using the same labels identified by the original authors ^28^. To this end, we computed the mean count of all cells in each cell-type separately. We then performed eQTL mapping as we have done extensively in the past ^27, 29^, by appropriate normalization and standardization of the data, and false discovery test correction (Methods). We found that alignment generally improved the number of eQTL detected, but only slightly (2–10%, Figure 5b).

Owing to the pooled design employed by the authors, the scRNA-seq data showed little to no batch effects which can explain the subtle improvement of eQTL detection power after alignment. To test this possibility, we next added batch effects (Methods) to each of the batches to simulate a more practical data collection pipeline, wherein clinical samples are collected continuously and immediately processed for scRNA-seq. UMAP of the unaligned data clearly shows the introduced batch effect (Figure 5c), which is consistent with scRNA-seq data from clinical samples that were immediately processed after collection (Figure 2a). When using unaligned data with simulated batch effects to identify eQTLs, the number of eQTLs identified in each cell-type was reduced by an average of 40–60%. These observations demonstrate the necessity to account for potential batch effects in eQTL mapping studies that use scRNA-seq data. Thus, we next identified eQTLs starting from data aligned using different methods. We found that eQTL mapping starting with data aligned using `Dmatch` resulted in the most eQTLs identified (1.3–1.5 fold as many as starting with unaligned data) compared to data aligned using other methods (0.9–1.1 fold as many) at 10% FDR (Figure 5d). To visualize the differences in eQTLs recovered from data aligned using the different methods, we compared the significance of the association for the QTLs. We found that the linear regression *p*-values were generally highly correlated, but in many cases were more significant for `Dmatch`-aligned data compared to data aligned using other methods (Supplementary Figure 11). This suggests that data aligned using `Dmatch` improves QTL mapping power. We also validated the eQTLs mapped using data from the DICE consortium, which collected population-level bulk RNA-seq data for a large number of sorted cell-types. We found that while the rates at which eQTLs identified from data aligned using the different methods were largely similar, `Dmatch` identified more eQTLs, which generally led to a higher number of eQTLs that are replicated overall (Figure 5e and Supplementary Figure 12).

Thus, we conclude that alignment of scRNA-seq data increase eQTL mapping power. However, the improvement in mapping power varies across alignment methods and is likely to be reduced when biological variation is removed due to over-correction.

## Discussion

There has been a great interest in using scRNA-seq to discover gene regulatory signatures that underlie various biological processes and phenomena. One major obstacle to fully exploit the power of scRNA-seq is the extensive batch and technical effects in scRNA-seq experiments. These unwanted effects not only limit the re-usability of scRNA-seq data produced by different groups, but makes it difficult to compare two scRNA-seq experiments generated by the same group and individual. To overcome this obstacle, a growing number of methods were proposed to “align” scRNA-seq experiments to facilitate comparison and allow downstream analysis of the merged datasets. Many of these alignment strategies have been designed to align datasets that have been generated using different scRNA-seq platforms or even to align scRNA-seq data from different species. However, there is a paucity of studies that evaluate the performance of these methods on aligning datasets that were generated using the same methods in perhaps different contexts, e.g. healthy versus disease condition. Indeed, while most methods appear to perform well on the task of aligning conspicuous cell clusters together (e.g. the monocytes from one dataset to monocytes of another dataset), it is unclear whether more subtle biological variation is lost during the alignment process.

We developed `Dmatch` to align scRNA-seq experimental data that are obtained using the same scRNA-seq platform to allow downstream analyses including differential gene expression analysis. `Dmatch` uses an external panel of reference transcriptomes (Primary Cell Atlas ^13^) that was compiled from over 100 separate studies to identify anchor cell-types that can be used for identifying batch or technical effects. While the cell-type annotation in the Primary Cell Atlas may be relatively coarse (95 distinct cell-type annotations), we found that `Dmatch` was able to find appropriate anchors to align samples consisting of immune cell-types. Although we expect this strategy to work less well for more specialized scRNA-seq samples that contain cell-types that are not represented in the Primary Cell Atlas, we envisage that ongoing efforts, such as that of the Human Cell Atlas ^30^, will establish better cell-type references that can be used by `Dmatch` to identify more appropriate anchor cell-types. `Dmatch` uses kernel density matching to estimate two parameters, a translation parameter and a rotation parameter, to align two scRNA-seq datasets. Because `Dmatch` transforms data linearly and this transformation depends only on two parameters, we reasoned that the resulting transformed data should be less prone to over-correction. The resistance to over-correction, however, comes at the cost of not being able to detect, and therefore correct for, batch or technical effects that are entirely nonlinear. Indeed, we found that alignments from `Dmatch` resemble uncorrected data more so than compared to alignments from other methods, suggesting that the corrections made by `Dmatch` are relatively conservative. Nevertheless, `Dmatch` performed similarly or favorably to all evaluated methods in terms of reduction of batch effects in simulated data and in terms of producing correctly aligned cell-type clusters across samples by visual inspection of PCA or UMAP plots.

A major difference we observed between the evaluated alignment methods in this study was that `Dmatch` appeared to be less or not prone to over-correction, while Harmony, scMerge, Seurat V3, and to some extent, MNN, all showed signs of over-correction. For example, we found several cell clusters that likely represented distinct cell states (e.g. pro-inflammatory and normal monocyte), but were made indistinguishable in alignments obtained using Harmony, scMerge, and Seurat V3. Even in the case of MNN, the differences in gene expression that distinguished these clusters were often reduced substantially. Thus, while all methods appear to work relatively well in terms of correcting batch or technical variation across scRNA-seq samples, methods vary dramatically in terms of their propensity to remove true biological variation. Indeed, we found that data aligned using scMerge showed the strongest signature of over-correction in data that we collected. In this study, we also found that over-correction can impact eQTL mapping. Indeed, while eQTL analysis is generally tolerant to under-correction, as covariates can be added to the linear regressions that test for associations, over-correction of true biological – in this case inter-individual – variation will strictly reduce eQTL mapping power.

In summary, we present `Dmatch`, a novel method for alignment of scRNA-seq data, which allows users to integrate multiple scRNA-seq experiments for downstream analyses. While we found that all evaluated methods performed well in terms of their ability to remove batch and technical effects, only `Dmatch` alignments appeared to be resistant to over-correction. Thus, `Dmatch` unlocks several important applications such as differential gene expression analysis and eQTL mapping using scRNA-seq data. These applications are becoming staple tools in genomic studies as scRNA-seq continues to increase in popularity and maturity.

## Methods

### Identifying anchor cells by projecting scRNAseq to a reference panel

We propose to use reference transcriptomes from the Primary Cell Atlas, a meta-analysis of a large number of publicly available microarray datasets compiled from human primary cells (745 samples, from over 100 separate studies). After pre-processing using quantile normalization and feature selection, we obtained a panel of 5,209 gene measurements for 95 annotated cell-types (e.g. epithelial cells, fibroblasts and endothelial cells, etc…) for projection. To identify subpopulations from the observed single cells, we projected all cells from our scRNA-seq samples to this reference panel by quantifying the Pearson or Spearman Correlation between measurements for each single cell and the measurements for the 95 annotated cell-types. We then enforced sparsity to the original Pearson correlation matrix, kept only the top five primary cell lines in the reference panel which were highly correlated with any cell in the experimental data sets and set the rest to be zero, and kept only **the primary cell lines in the reference panel which were among the top five primary cells lines of more than twenty cells in the experimental data sets and set the rest to be zero**. The cut-offs were selected empirically for 10X data and could be adjusted based on the quality of signals in the datasets. Of note, the identities of the single cells were described by a 95-dimensional projected vector on the reference panel. We further performed a bi-clustering which allowed simultaneous clustering of the rows and columns on the sparsity-enforced Pearson correlation matrix. We used hlcust() implemented in R for the hierarchical clustering with Euclidean distance as the distance metric. In our experiments, complete linkage, Ward’s minimum variance method, and Ward’s minimum variance square linkage method achieved the best performance. Finally, we determined clustering membership based on the bi-clustering results. The cell clusters shared across samples were used as anchor cell-types for alignment. If two samples share more than two cell clusters, the default is to select two clusters based on 1) the number of cells in each cluster: a recommended cluster has at least 5% of the total number of cells in each sample and 2) the results from a Shapiro test on the normality.

### Removal of batch effects

We assume the batch effects can be represented as linear transformations on reduced dimensions. We used PCA for dimension reduction in the analysis, which could be replaced by other non-linear alternatives. We assume that the density of each selected anchor cell-type on a pair of PCs follows a 2-D Gaussian distribution. Taking advantage of orthogonality, we can perform the correction independently on every consecutive pair of top PCs. In the default setting, `Dmatch` accounts for variation loaded on top 30 PCs. Overall, we seek to correct for batch effects through applying linear transformations estimated from density matching.

To illustrate the general ideas, we describe the alignment procedure for measurements from two cell-types, type A and type B cells in the individual 1 and 2, respectively. At population level, the density for measurements in the individual 1 is *g* and the density for measurements in the individual 2 is *p* without any perturbation. Let *Y*′ represent measurements of type A cells from individual 1, *X*′ represent measurements of type B cells from individual 1, *Y* represent measurements of type A cells from individual 2, *X* represent measurements of type B cells from individual 2 after some unknown affine perturbation on the original density. Let *D*(*q, p*) represent some distance measure for two densities, for example, Hellinger distance, total variation distance, Chi-square distance or Kullback-Leibler divergence. We seek an affine transformation on *{Y, X}*, including a translation *d* and a linear map *A*, which can minimize

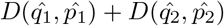

where 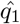 and 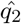 are the estimated density functions based on *Y*′ and *X*′, respectively; 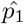 and 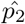 are the estimated density functions based on *Y A − d* and *X A − d*, respectively. The solution of this problem depends on the distance measure as well as the density estimation method. We use Kullback-Leibler (KL) divergence, which possesses good properties for our application and results in a closed-form objective. The KL divergence from *p* to *q* is defined as:

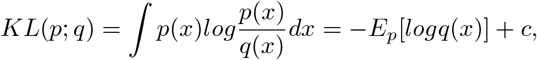

where q is reference density that we want to align to. It is a measure of how much “evidence” each sample will on average bring when we are trying to distinguish *p*(*x*) from *q*(*x*) and when we are sampling from p(x). Accordingly, our problem can be formulated as:

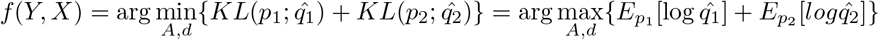

This is equivalent to calculate maximum likelihood estimators for *A, d* given empirical density of *q*_1_ and *q*_2_. We use Gaussian distributions to estimate the densities for *X*, *Y*, *X*′, and *Y*′, respectively. Let ***μ***_*y*_, **D**_1_ and ***μ***_*x*_, **D**_2_ represent the estimated mean vector and covariance matrix for **Y**′ and **X**′, respectively. The objective becomes the following;

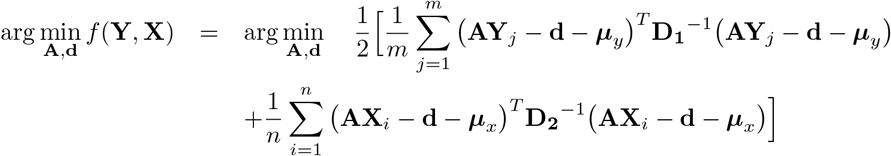

The gradients respect to **A** and **d** equal to the following:

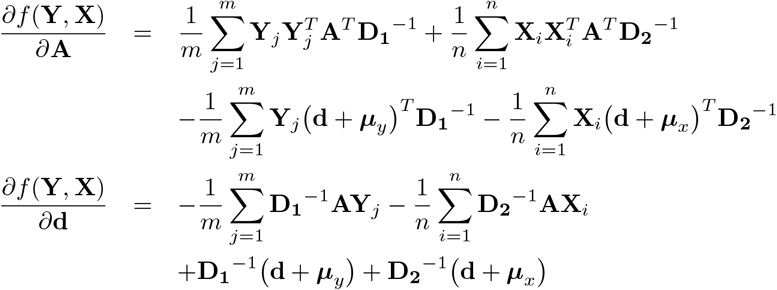

Further simplification leads to the following matrix form representation:

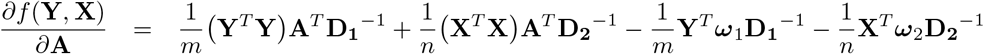

where *ω*_1_ = 1_*m*_ ⊗ (d + *μ*_*y*_)^*T*^ and *ω*_2_ = 1_*n*_ ⊗ (d + *μ*_*x*_)^*T*^.

We use coordinate descent to solve the above estimation problem. At each step, **d** will be updated by:

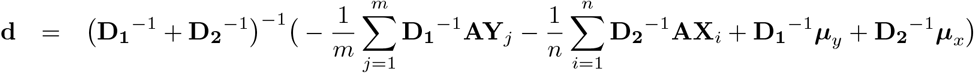

Theoretically, *A* is allowed to take many forms of linear transformation. In the default implementation, we constrained the linear map in the affine transformation to a single rotation, dramatically reducing the search space of parameters, i.e.,

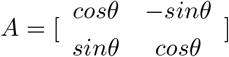

As such, we eplace the gradient search by angle search in the algorithm:

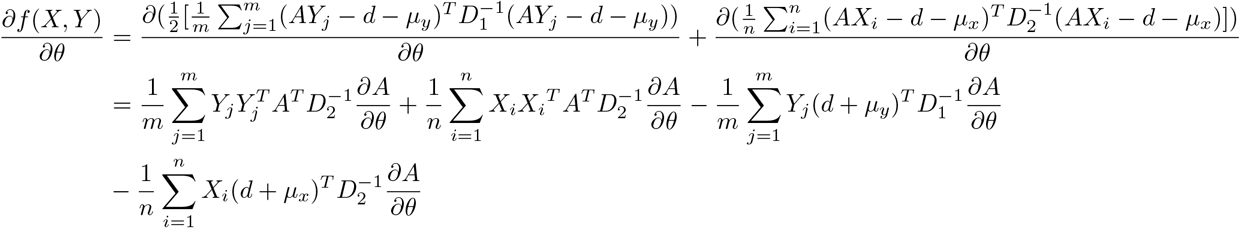

where

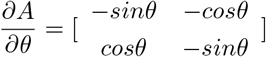

### Simulation study

We adopt the simulation model from MNN paper to evaluate the performance of alignment methods. Case 1 (easy case): We considered a four-component Gaussian mixture model in 2-D (to represent the low dimensional biological subspace), where each mixture component represents a different simulated cell-type. In other words, we generate four Gaussian distributions with different means and standard deviations to imitate four cell-types.

Two data sets with N = 1000 cells were drawn with equal mixing coefficients (roughly 0.25/0.25/0.25/0.25 for the two batches) for the four cell-types. **Two batches share all their four cell-types and have evenly distributed cellular proportions.**

We then projected both data sets to G = 100 dimensions using the same random Gaussian matrix, to simulate observed high-dimensional gene expression data. Batch effects were incorporated by generating a Gaussian random vector for each data set and adding it to the expression profiles for all cells in that data set.

Case 2 (difficult case): Compared to the easy case, we simulate six Gaussian distributions in 2-D space. The first batch has the first four Gaussian distributions as its four cell-types, and the second batch has the last four Gaussian distributions as its four cell-types. Therefore, those two batches partially share cell-type 3 and 4 (or cluster 3 and 4). Of note,two batches share two cell-types with each other (partially sharing): cluster 3 and cluster 4, and the proportion of each cell-type in each batch is approximately even (0.25/0.25/0.25/0.25).

### Consistency of using different cell clusters as anchors

To evaluate the consistency of using different cell clusters as anchors for `Dmatch`, we selected HC1 PBMC and SLE PBMC as test samples and aimed to demonstrate that any combinations of clusters identified using `Dmatch` give consistent alignments. For the alignment between HC and SLE PBMCs, `Dmatch` identified 3 possible alignment clusters (2, 3, and 4). This allows `Dmatch` to use as anchors a combination of cluster 2 and 3, cluster 2 and 4, cluster 3 and 4, and cluster 2 and 3 and 4. As a baseline, we compared the alignments using the 4 combinations of clusters against alignments of SLE PBMC on RA1 PBMC (baseline 1) and of SLE PBMC on SLE BMMC (baseline 2), which are expected to produce different alignments.

To quantitatively measure how different the alignments were, we calculated the mean squared error (MSE) distance across alignments (on the top 30 PCs) using different anchors and the baseline comparisons. We found that the alignments using different combinations of selected clusters have much smaller MSE than compared to the two baseline alignments (see tables 3 and 4 below), supporting the consistency of using different cell clusters as anchors.

**Table 1:**
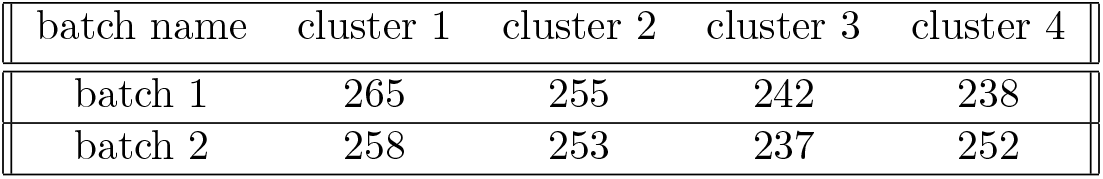
Cell numbers for each cell-type cluster in the two simulated batches for the easy case.

**Table 2:**
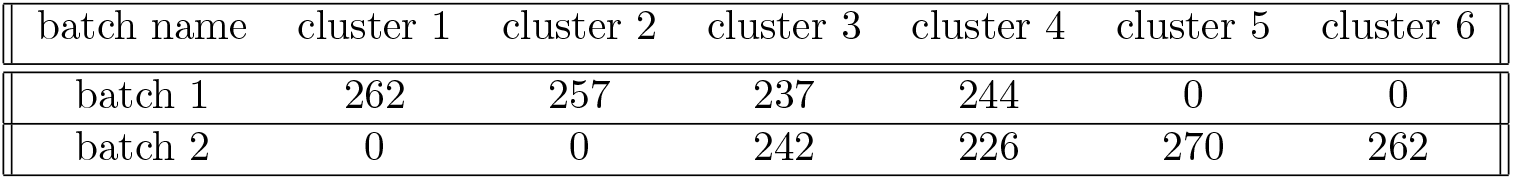
Cell numbers for each cell-type cluster in the two simulated batches for the difficult case.

| batch name | cluster 1 | cluster 2 | cluster 3 | cluster 4 | cluster 5 | cluster 6 |
| --- | --- | --- | --- | --- | --- | --- |
| batch 1 | 262 | 257 | 237 | 244 | 0 | 0 |
| batch 2 | 0 | 0 | 242 | 226 | 270 | 262 |

**Table 3:**
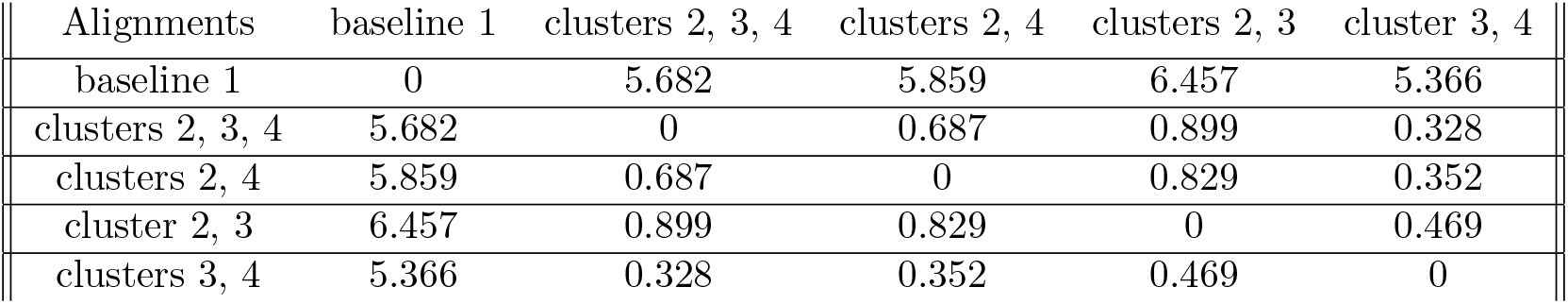
MSE (top 30 PCs) of data after alignment using different anchors plus baseline 1.

| Alignments | baseline 1 | clusters 2, 3, 4 | clusters 2, 4 | clusters 2, 3 | cluster 3, 4 |
| --- | --- | --- | --- | --- | --- |
| baseline 1 | 0 | 5.682 | 5.859 | 6.457 | 5.366 |
| clusters 2, 3, 4 | 5.682 | 0 | 0.687 | 0.899 | 0.328 |
| clusters 2, 4 | 5.859 | 0.687 | 0 | 0.829 | 0.352 |
| cluster 2, 3 | 6.457 | 0.899 | 0.829 | 0 | 0.469 |
| clusters 3, 4 | 5.366 | 0.328 | 0.352 | 0.469 | 0 |

**Table 4:** MSE (top 30 PCs) of data after alignment using different anchors plus baseline 2.

| Alignments | baseline 2 | clusters 2, 3, 4 | clusters 2, 4 | clusters 2, 3 | clusters 3, 4 |
| --- | --- | --- | --- | --- | --- |
| baseline 2 | 0 | 13.729 | 13.290 | 14.244 | 13.322 |
| clusters 2, 3, 4 | 13.729 | 0 | 0.687 | 0.899 | 0.328 |
| clusters 2, 4 | 13.290 | 0.687 | 0 | 0.829 | 0.352 |
| clusters 2, 3 | 14.244 | 0.899 | 0.829 | 0 | 0.469 |
| clusters 3, 4 | 13.322 | 0.328 | 0.352 | 0.469 | 0 |

## Performance evaluation

In addition to the visual inspection of PC plots, we computed the silhouette coefficients and Random Adjusted Index to evaluate the performance of different methods on simulated data.

### Silhouette Coefficients

Let a(i) be the average distance of cell i to all other cells within the same cluster as i and b(i) be the average distance of cell i to all cells assigned to the neighboring cluster, i.e., the cluster with the lowest average distance to the cluster of i. The original silhouette coefficient for cell i is defined as:

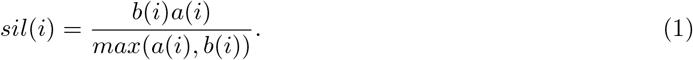

When b(i) is much greater than a(i), the silhouette coefficient approaches to 1. On the contrary, when a(i) is much greater than b(i), the silhouette coefficient approaches to −1. Ideally, we would want ¡b(i)¿ to be relatively way larger than ¡a(i)¿ so that silhouette coefficient will approach to 1. Finally, we take the arithmetic average of all cells’ silhouette coefficients in a batch to get a single number within [−1,1] to assess different batch correction methods.

### Adjusted Random Index (ARI)

We used adjusted Rand index (ARI) to evaluate the concordance of clustering results with respect to the true cell-type labels, or the match between true cell labels and the assigned cell labels after performing clustering analysis on corrected data. Considering the cells of corrected data are partitioned into different classes with respect to cell-type labels, let a be the number of pairs of cells partitioned into the same class by a clustering method and in fact belong to that same class; b be the number of pairs of cells partitioned into the same cluster but in fact belong to different classes; c be the number of pairs of cells partitioned into different clusters but belong to the same class; and d be the number of pairs of cells from different classes partitioned into different clusters. Then the ARI is calculated as

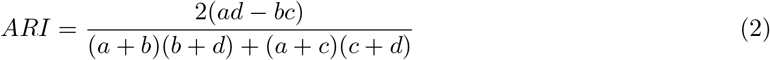

A high ARI approaching 1 indicates a high concordance between the clustering result and known cell-type information, and thus good performance of the method for correcting batch effects.

### Data collection

Samples of peripheral blood (PB) and bone marrow (BM) were obtained from three RA patients, one AS patients, one SLE patients, and one HC. And synovial fuid (SF) samples were obtained from the knee joints of two RA patients. RA, AS or SLE patients fulfilled the 2010 American College of Rheumatology /European League Against Rheumatism (ACR/EULAR) RA classification criteria ^31^, the SpondyloArthritis International Society classification criteria for axial spondylarthritis ^32^, or the ACR Revised Criteria for Classification of SLE ^33^, respectively. Donors were enrolled at Department of Clinical Immunology of Xijing Hospital in Xi’an, and their clinical characteristics are summarized in Supplementary Table 1. Donors with chronic disease, cancer and chronic infections such as Hepatitis B, C, and HIV were excluded. PB, BM, and SF were collected by EDTA treated Vacutainer tubes in a sterile manner (BD Biosciences, Franklin Lakes, NJ, USA). SF was diluted with two volumes of phosphate-buffered saline (PBS) and treated with bovine testicular hyaluronidase (10mg/ml; Sigma-Aldrich, St Louis, MO, USA) for 30 min at 37C. Mononuclear cells were isolated using Ficoll-Paque gradient separation (GE Healthcare Bio-Sciences, Pittsburgh, PA, USA). This research was approved by the ethical standards committee of Xijing Hospital (KY20192006-C-1). All donors provided written informed consent in accordance with the Declaration of Helsinki.

### Identification of cell clusters using Seurat and markers

We used Seurat (version 2.3) with default recommended settings on each of our scRNA-seq samples separately to identify clusters (referred to as Seurat clusters). To annotate Seurat clusters, we inspected the top 10 marker genes that were the most over-expressed in each cluster compared to the remaining cells as identified using the “FindAllMarkers” function from Seurat. More specifically, we used 2–3 representative markers to annotate the following cell-types CD4+ T cells (*CD3D*, *IL7R*), CD8+ T cells (*CD3D, CD8A, CD8B*), NK cells (*NKG7,GNLY*), B cells (*MS4A1, CD79A*), CD14+ Monocyte (*LYZ, CD14*), CD16+ Monocyte (*LYZ, FCGR3A*), Red Blood Cells (*HBB, HBA1, HBA2*), Dendritic Cells (*FCER1A, CST3*).

### Alignment of scRNA-seq data from patient and healthy biopsies

To align the scRNA-seq data from patient and healthy individual biopsies (PBMC, PBMC+BMMC, and PBMC+SF), we used the default parameters for Seurat V3 ^17^, MNN (scran 1.9.39) ^11^, scMerge ^18^(0.1.9.1), and Harmony ^19^ (0.0.0.9000) and used the 30 first PCs as input data (except for scMerge, for which the full dataset was used as input, as required).

`Dmatch` aligns scRNA-seq samples by finding the alignment parameters that minimize the difference in the densities between anchor cell clusters from the “target” and “source” scRNA-seq samples and by applying the transformation to all cells including cells in the “source” dataset. Thus, the alignment process can fail if batch effects are highly inconsistent among cell clusters (and thus anchor cell clusters). To detect such inconsistencies, `Dmatch` computes an alignment vector for each anchor cell clusters as the linear transposition in a 2D PC space from the center of an anchor cell cluster from the “source” sample to the same anchor cluster of the “target” sample. If the linear transpositions of two anchor cell clusters are very different, then the batch effects are likely to be highly inconsistent and result in the incorrect estimation of alignment parameters. In other words, differences in the linear transposition are indicative that different anchor cell clusters provide conflicting information regarding how batch effect should be corrected.

We found no inconsistent batch effects for our PBMC and PBMC+BMMC alignments, and therefore used the default parameters for `Dmatch` to align PBMC and PBMC+BMMC samples with 30 PCs as input. However, when we computed the angle between the alignment vectors for the alignment of RA1 SF onto RA1 PBMC, we found that the angles were small (consistent) for pairs of anchor clusters in PCs 1–6, but large (inconsistent batch effects) for many pairs of anchor clusters in PCs 7 and onwards (compare Table 5 to Table 6). Thus, we used the 6 first PCs as input for our PBMC+SF alignment. We found no qualitative difference when we used the first 6 PCs or all 30 PCs for the other methods.

**Table 5:**
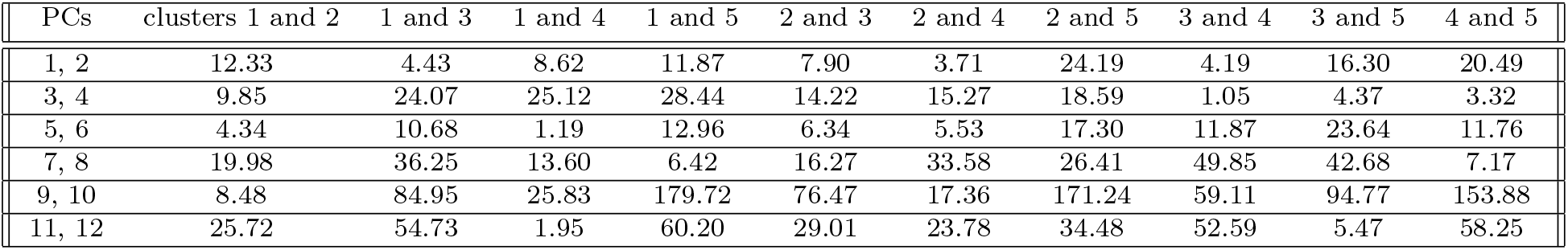
Angles between the alignment vectors for pairs of clusters for the SLE PBMC to HC SLE PBMC alignment. Angles are relatively small, which indicate consistent batch effect correction across anchor cell clusters.

| PCs | clusters 1 and 2 | 1 and 3 | 1 and 4 | 1 and 5 | 2 and 3 | 2 and 4 | 2 and 5 | 3 and 4 | 3 and 5 | 4 and 5 |
| --- | --- | --- | --- | --- | --- | --- | --- | --- | --- | --- |
| 1, 2 | 12.33 | 4.43 | 8.62 | 11.87 | 7.90 | 3.71 | 24.19 | 4.19 | 16.30 | 20.49 |
| 3, 4 | 9.85 | 24.07 | 25.12 | 28.44 | 14.22 | 15.27 | 18.59 | 1.05 | 4.37 | 3.32 |
| 5, 6 | 4.34 | 10.68 | 1.19 | 12.96 | 6.34 | 5.53 | 17.30 | 11.87 | 23.64 | 11.76 |
| 7, 8 | 19.98 | 36.25 | 13.60 | 6.42 | 16.27 | 33.58 | 26.41 | 49.85 | 42.68 | 7.17 |
| 9, 10 | 8.48 | 84.95 | 25.83 | 179.72 | 76.47 | 17.36 | 171.24 | 59.11 | 94.77 | 153.88 |
| 11, 12 | 25.72 | 54.73 | 1.95 | 60.20 | 29.01 | 23.78 | 34.48 | 52.59 | 5.47 | 58.25 |

**Table 6:** Angles between the alignment vectors for pairs of clusters for the RA1 SF to HC PBMC alignment. Angles for PCs 7–12 are large, which indicate inconsistent batch effect correction across many anchor cell clusters.

| PCs | cluster1 and 2 | 1 and 3 | 1 and 4 | 2 and 3 | 2 and 4 | 3 and 4 |
| --- | --- | --- | --- | --- | --- | --- |
| 1, 2 | 3.97 | 14.56 | 12.69 | 10.59 | 8.71 | 1.88 |
| 3, 4 | 58.12 | 33.13 | 41.44 | 24.99 | 16.68 | 8.31 |
| 5, 6 | 2.77 | 12.49 | 31.85 | 9.72 | 29.07 | 19.36 |
| 7, 8 | 10.82 | 70.35 | 100.82 | 59.53 | 111.64 | 171.18 |
| 9, 10 | 27.56 | 179.07 | 146.65 | 153.37 | 174.21 | 32.42 |
| 11, 12 | 48.80 | 146.48 | 163.28 | 97.68 | 114.48 | 16.80 |

Functions for computing alignment vectors and their angles are included in the `Dmatch` R package and can be used to select the appropriate number of PCs to input when the batch effect vectors are inconsistent.

### Differential expression analysis

To estimate differential expression across Seurat clusters, we used limma-trend as implemented in ^34^ and in which limma-trend was shown to perform well in nearly all major evaluation criteria including true positive rates and maximum false positive rates. To evaluate the differences between differential expressed genes (DEGs) identified using unaligned data and data aligned using the various methods (Figs 3a,b), limma-trend was applied to identify DEGs across all pairs of Seurat clusters from all samples combined. To be considered a DEG, we required a p-value smaller than 10^−10^ and an absolute log_2_ fold change difference to be greater than 1. Although these cutoffs were chosen arbitrarily, we found that varying the p-value cutoff to 10^−5^, 10^−20^ or the absolute log_2_ fold change difference cutoff to 0.5 or 1.5 did not qualitatively change our interpretations.

To identify DEG that are shared or specific to a method (Figure 3a and b), we simply counted the number of DEG that were identified at the aforementioned p-value and log_2_ fold change cutoffs for each of the different tools. We then plotted the number of DEG that were identified specifically using method A (positive y-axis), specifically using method B (negative y-axis), and DEG that were identified using both methods (positive x-axis) for all pairwise comparisons across Seurat clusters. Comparisons across Seurat clusters from the same scRNA-seq sample were colored in red, and comparisons across Seurat clusters with the same cell-type assignment were marked as crosses (again, see Figure 3a and b).

To produce the heatmaps in Figure 4a, we again called DEG across Seurat cluster across PBMC and BMMC scRNA-seq samples. We counted the number of DEGs for each pairwise comparison and used the `heatmap.2` command from the R package `gplots` to produce the heatmap using hierarchical clustering with the average linkage clustering.

### Population PBMC scRNA-seq dataset and simulated batch effects

We analyzed processed data from ^28^ and used their cell-type annotations. When performing UMAP on processed data, we observed that samples mixed well, indicative of limited batch effects. To add batch effects to the scRNA-seq data in order to simulate a more simple collection procedure without the need to pool samples together, we once again used the simulation model from MNN ^11^. Briefly, perturbation effects were incorporated by generating a Gaussian random vector for each data set and adding it to the expression profiles for all cells in that dataset. More specifically, gene expression profiles were LogNormalized, and eight perturbation parameters were generated for each of the eight samples from ^28^ (mean, standard deviation of the Gaussian random vector): list(c(1,0.28),c(1.6,0.4), c(1.8,0.35), c(2.5,0.45), c(2.2,0.38), c(1.8,0.22), c(1.95,0.4), c(2.0,0.3)). These eight Gaussian random vectors (each with length: *n* cells *× n* genes) were then added to each of the eight samples. The above parameters were chosen based on the PCA and tsne plots.

### eQTL mapping

To map eQTL in scRNA-seq data, we computed, for all individuals separately, the mean expression levels of all cells in each annotated cell-types: B cells, CD4+ T cells, CD8+ T cells, Dendritic cells, NK cells, Monocytes, and combined cells (pseudo-bulk, or bulk). We then mapped eQTL in each cell-type separately using a standard pipeline eQTL pipeline by quantile normalizing expression levels across genes within individuals and standardize the expression across individuals ^27^. We then used `fastqtl` ^35^ to identify associations between genetic variants and gene expression levels and the Benjamini-Hochberg procedure to control false discovery rates at the 0.05, 0.10, and 0.20 levels. No PCs were used as covariates.

### Replication using DICE eQTLs

To replicate the eQTLs identified using the population scRNA-seq data, we quantified gene expression levels from DICE RNA-seq data for each different cell-type separately using Kallisto vx ^36^. We then used standard eQTL pipeline ^27^ and `fastqtl` ^35^ with 3 genotype PCs and a variable number of phenotypic PCs. More specifically, PCA on genotype data was performed using the smartpca program from EIGENSOFT version 6.1.4, after pruning SNPs using PLINK v1.9 with parameters --`indep-pairwise 50 2 0.2`. Individual outliers reported by smartpca were excluded from eQTL mapping. Phenotypic PCs were calculated in R using the `prcomp` function. We empirically chose the number of phenotypic PCs that maximized the number of significant eQTLs (5% FDR) in each cell-type (Supplementary Table 2).

To determine the replication of a eQTL from aligned scRNA-seq data, we asked whether the linear regression between the gene expression and the most significantly associated SNP in the aligned scRNA-seq data was also significant in each of the cell-types in the DICE dataset (*p* < 0.05). To determine the replication in “any” cell-type, we asked whether the linear regression was significant, correcting for the number of cell-types 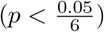.

## Acknowledgement

We thank Sebastian Pott for comments on the manuscript. This work was supported by the US National Institutes of Health (R01GM130738 to Y.I.L, and GM126553 to M.C.) and by the National Basic Research Program of China (NO.2015CB553704) to P.Z.. M.C. also acknowledges support from the Sloan Research Foundation Fellowship and a HCA Seed Network grant from the Chan Zuckerberg Initiative. This work was completed in part with resources provided by the University of Chicago Research Computing Center.

## Author Contributions

Y.I.L. and M.C. conceived of the project. M.C., Q.Z. performed analyses and implemented the software. Y.I.L., Z.M, and L.W. also performed analyses. Y.I.L. and M.C. wrote the manuscript with contribution from Q.Z.. Z.P. collected the experimental data.

## References

[1] Kumar, R. M. et al. Deconstructing transcriptional heterogeneity in pluripotent stem cells. Nature 516, 56–61 (2014).

[2] Villani, A. C. et al. Single-cell RNA-seq reveals new types of human blood dendritic cells, monocytes, and progenitors. Science 356(2017).

[3] Durruthy-Durruthy, R. et al. Reconstruction of the mouse otocyst and early neuroblast lineage at single-cell resolution. Cell 157, 964–978 (2014).

[4] Bendall, S. C. et al. Single-cell trajectory detection uncovers progression and regulatory coordination in human B cell development. Cell 157, 714–725 (2014).

[5] Shalek, A. K. et al. Single-cell RNA-seq reveals dynamic paracrine control of cellular variation. Nature 510, 363–369 (2014).

[6] Keren-Shaul, H. et al. A Unique Microglia Type Associated with Restricting Development of Alzheimer’s Disease. Cell 169, 1276–1290 (2017).

[7] Golumbeanu, M. et al. Single-Cell RNA-Seq Reveals Transcriptional Heterogeneity in Latent and Reactivated HIV-Infected Cells. Cell Rep 23, 942–950 (2018).

[8] Macosko, E. Z. et al. Highly Parallel Genome-wide Expression Profiling of Individual Cells Using Nanoliter Droplets. Cell 161, 1202–1214 (2015).

[9] Brennecke, P. et al. Accounting for technical noise in single-cell RNA-seq experiments. Nat. Methods 10, 1093–1095 (2013).

[10] Yuan, G. C. et al. Challenges and emerging directions in single-cell analysis. Genome Biol. 18, 84 (2017).

[11] Haghverdi, L., Lun, A. T. L., Morgan, M. D. & Marioni, J. C. Batch effects in single-cell RNA-sequencing data are corrected by matching mutual nearest neighbors. Nat. Biotechnol. 36, 421–427 (2018).

[12] Butler, A., Hoffman, P., Smibert, P., Papalexi, E. & Satija, R. Integrating single-cell transcriptomic data across different conditions, technologies, and species. Nat. Biotechnol. 36, 411–420 (2018).

[13] Mabbott, N. A., Baillie, J. K., Brown, H., Freeman, T. C. & Hume, D. A. An expression atlas of human primary cells: inference of gene function from coexpression networks. BMC Genomics 14, 632 (2013).

[14] Wu, C., Jin, X., Tsueng, G., Afrasiabi, C. & Su, A. I. Biogps: building your own mash-up of gene annotations and expression profiles. Nucleic acids research 44, D313–D316 (2015).

[15] Li, H. et al. Reference component analysis of single-cell transcriptomes elucidates cellular heterogeneity in human colorectal tumors. Nat. Genet. 49, 708–718 (2017).

[16] Li, S. et al. Detecting and correcting systematic variation in large-scale RNA sequencing data. Nat. Biotechnol. 32, 888–895 (2014).

[17] Stuart, T. et al. Comprehensive integration of single-cell data. Cell (2019).

[18] Lin, Y. et al. scmerge: Integration of multiple single-cell transcriptomics datasets leveraging stable expression and pseudo-replication. bioRxiv (2018). arXiv:https://www.biorxiv.org/content/early/2018/09/12/393280.full.pdf.

[19] Korsunsky, I. et al. Fast, sensitive, and accurate integration of single cell data with harmony. bioRxiv (2018). arXiv:https://www.biorxiv.org/content/early/2018/11/05/461954.full.pdf.

[20] Rousseeuw, P. J. Silhouettes: A graphical aid to the interpretation and validation of cluster analysis. Journal of Computational and Applied Mathematics 20, 53–65 (1987).

[21] Santos, J. M. & Embrechts, M. On the use of the adjusted rand index as a metric for evaluating supervised classification. In Proceedings of the 19th International Conference on Artificial Neural Networks: Part II, ICANN ’09, 175–184 (Springer-Verlag, Berlin, Heidelberg, 2009).

[22] Kay, J. & Calabrese, L. The role of interleukin-1 in the pathogenesis of rheumatoid arthritis. Rheumatology (Oxford) 43 Suppl 3, iii2–iii9 (2004).

[23] Dinarello, C. A. Interleukin-1 in the pathogenesis and treatment of inflammatory diseases. Blood 117, 3720–3732 (2011).

[24] Wan, C. K., He, C., Sun, L., Egwuagu, C. E. & Leonard, W. J. Cutting Edge: IL-1 Receptor Signaling is Critical for the Development of Autoimmune Uveitis. J. Immunol. 196, 543–546 (2016).

[25] Law, C. W., Chen, Y., Shi, W. & Smyth, G. K. voom: Precision weights unlock linear model analysis tools for RNA-seq read counts. Genome Biol. 15, R29 (2014).

[26] Koch, A. E. Chemokines and their receptors in rheumatoid arthritis: future targets? Arthritis Rheum. 52, 710–721 (2005).

[27] Li, Y. I. et al. RNA splicing is a primary link between genetic variation and disease. Science 352, 600–604 (2016).

[28] van der Wijst, M. G. P. et al. Single-cell RNA sequencing identifies celltype-specific cis-eQTLs and co-expression QTLs. Nat. Genet. 50, 493–497 (2018).

[29] Li, Y. I. et al. Annotation-free quantification of rna splicing using leafcutter. Nature Genetics 50, 151–158 (2018).

[30] Regev, A. et al. The Human Cell Atlas. Elife 6(2017).

[31] Aletaha, D. et al. 2010 rheumatoid arthritis classification criteria: an American College of Rheumatology/European League Against Rheumatism collaborative initiative. Ann. Rheum. Dis. 69, 1580–1588 (2010).

[32] Rudwaleit, M. et al. The development of Assessment of SpondyloArthritis international Society classification criteria for axial spondyloarthritis (part II): validation and final selection. Ann. Rheum. Dis. 68, 777–783 (2009).

[33] Hochberg, M. C. Updating the American College of Rheumatology revised criteria for the classification of systemic lupus erythematosus. Arthritis Rheum. 40, 1725 (1997).

[34] Soneson, C. & Robinson, M. D. Bias, robustness and scalability in single-cell differential expression analysis. Nat. Methods 15, 255–261 (2018).

[35] Ongen, H., Buil, A., Brown, A. A., Dermitzakis, E. T. & Delaneau, O. Fast and efficient QTL mapper for thousands of molecular phenotypes. Bioinformatics 32, 1479–1485 (2016).

[36] Bray, N. L., Pimentel, H., Melsted, P. & Pachter, L. Near-optimal probabilistic RNA-seq quantification. Nat. Biotechnol. 34, 525–527 (2016).

