## Supplementary Figures for "Alignment of single-cell RNA-seq samples without over-correction using kernel density matching"

### Supplementary Figures for Alignment of single-cell RNA-seq samples without overcorrection using kernel density matching

Mengjie Chen[1,2*], Qi Zhan[3], Zepeng Mu[3], Lili Wang[3], Zhaohui Zheng[4,5], Jinlin Miao[4,5], Ping Zhu[4,5*], Yang I Li[1,2*]

[1] Section of Genetic Medicine, Department of Medicine, University of Chicago, IL

[2] Department of Human Genetics, University of Chicago, IL

[3] Committee on Genetics, Genomics & Systems Biology, University of Chicago, IL

[4] Department of Clinical Immunology, Xijing Hospital, Xi'an, China

[5] National Translational Science Center for Molecular Medicine, Xi'an, China

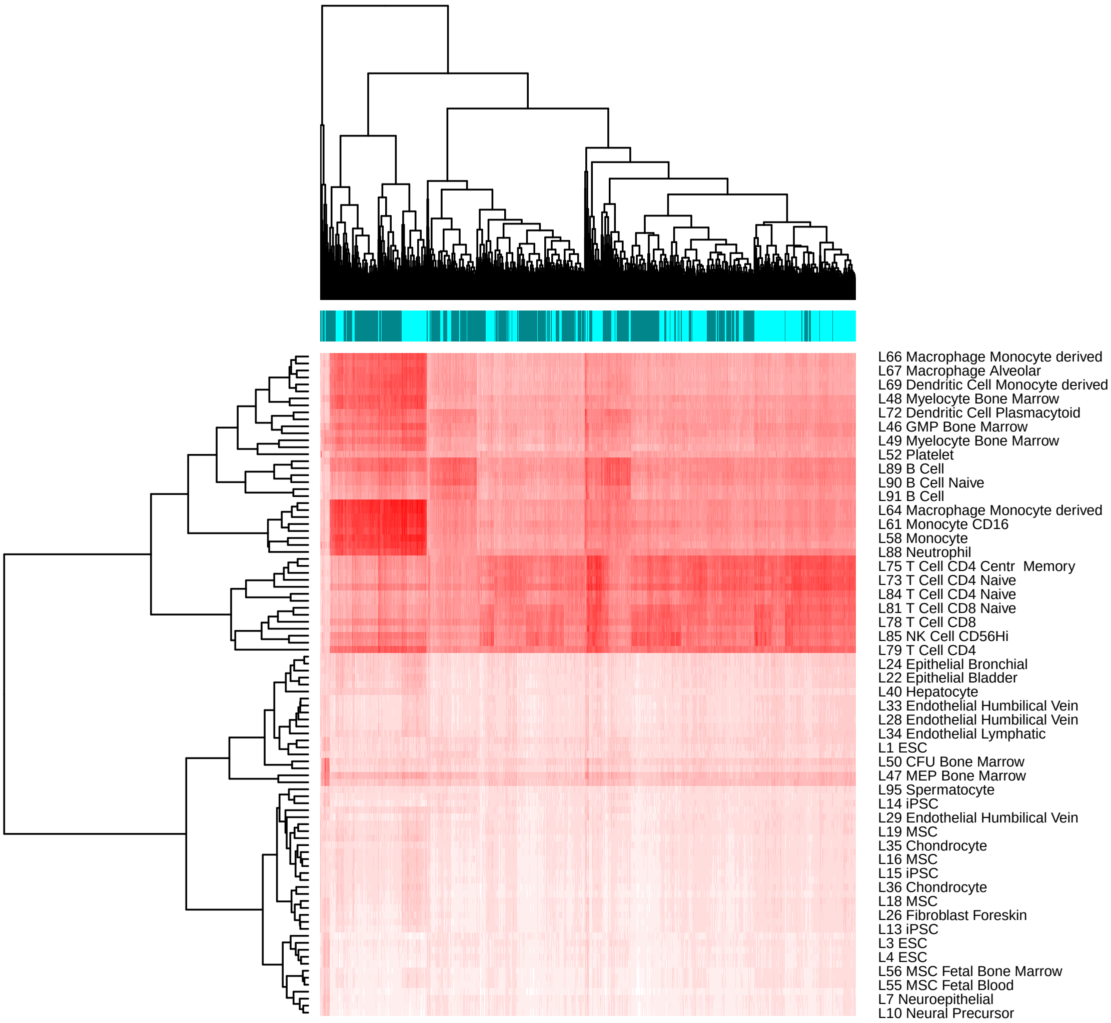


Supplementary Figure 1: `Dmatch` uses a large reference transcriptomes from the Primary Cell Atlas to identify subpopulations from the observed cells based on the projection. We found that the consistency of cell-type assignment was reduced if Pearson correlations of the 95-dimensional vectors were not set to zero except for those between the cell and the top five reference atlas cell-types with the highest Pearson correlation coefficient. When this sparsity was not induced, the biclustering becomes noisier and the anchors less apparent (compared to Fig. 1b).

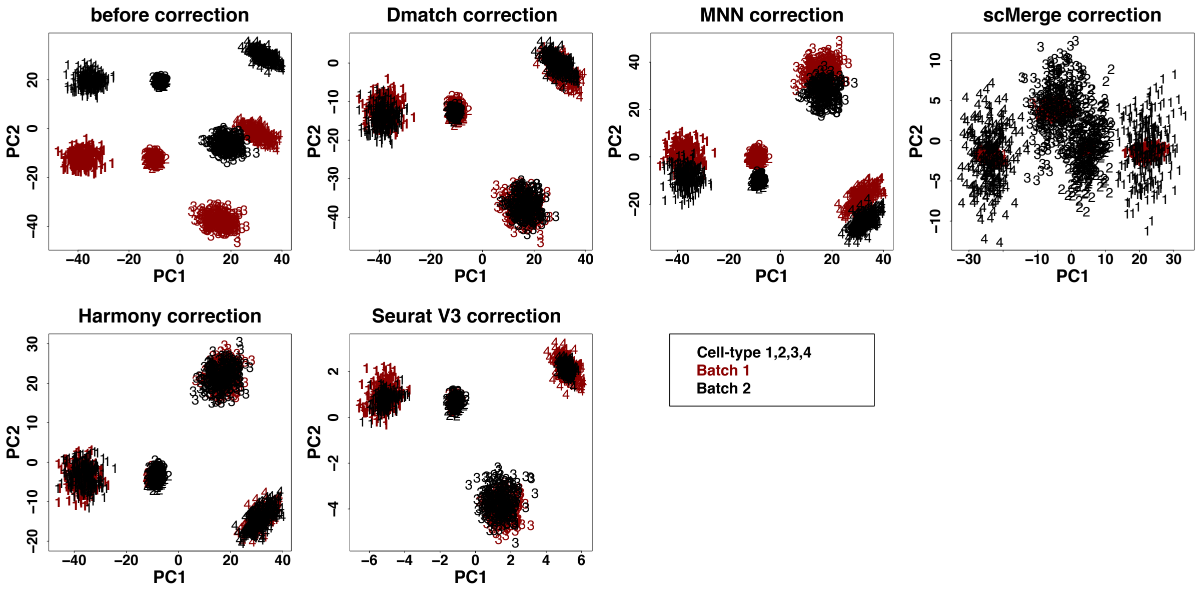


Supplementary Figure 2: Scatter plot of the first two PCs of unaligned and aligned simulated data using multiple alignment methods. Most methods were able to align samples in this easier case in which all cell-types were shared across the two samples/batches.

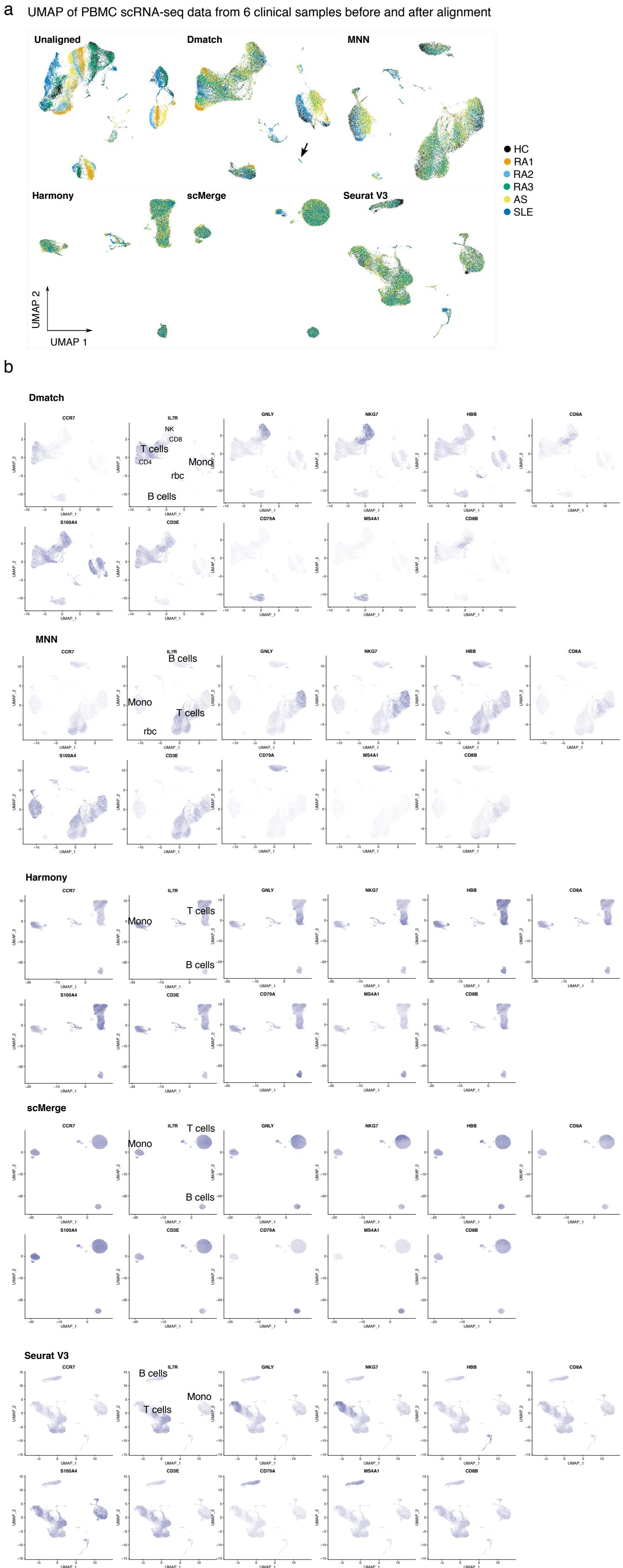


Supplementary Figure 3: **(a)** UMAP dimensionality reduction of PBMC scRNA-seq data from 6 clinical samples before and after alignment comparing `Dmatch`, MNN, Harmony, scMerge, and Seurat V3. **(b)** Monocytes, T cells and B cells are clearly separated in the UMAP clustering for all methods. Shades of purple represent marker expression in each cell relative to other cells in the same UMAP (arbitrary unit).

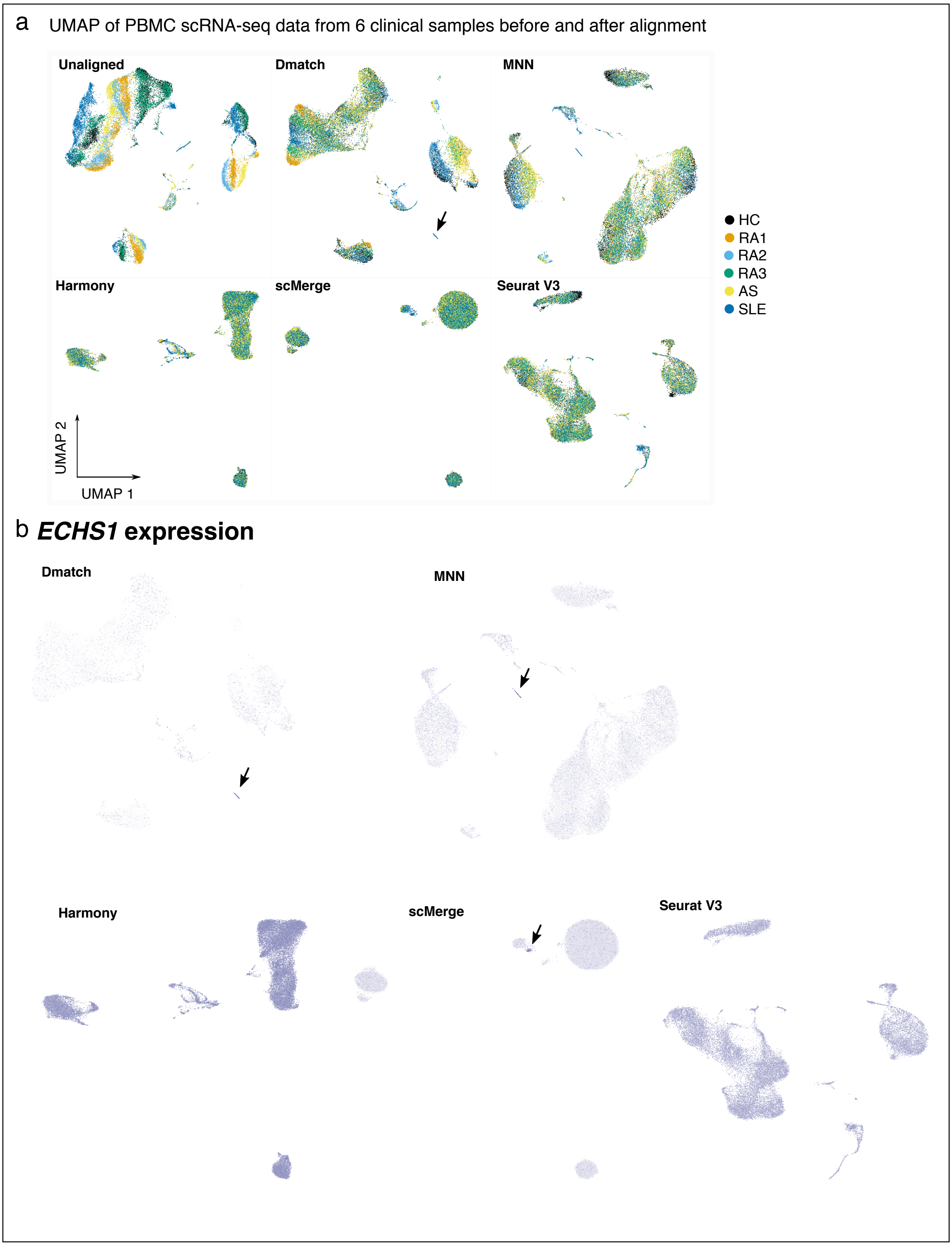


Supplementary Figure 4: **(a)** UMAP dimensionality reduction of PBMC scRNA-seq data from 6 clinical samples before and after alignment comparing `Dmatch`, MNN, Harmony, scMerge, and Seurat V3. **(b)** The marker *ECHS1* for the "kidney" cells cluster tags the SLE-specific cluster only in data aligned using `Dmatch`, MNN, and to some extent scMerge. Shades of purple represent marker expression in each cell relative to other cells in the same UMAP.

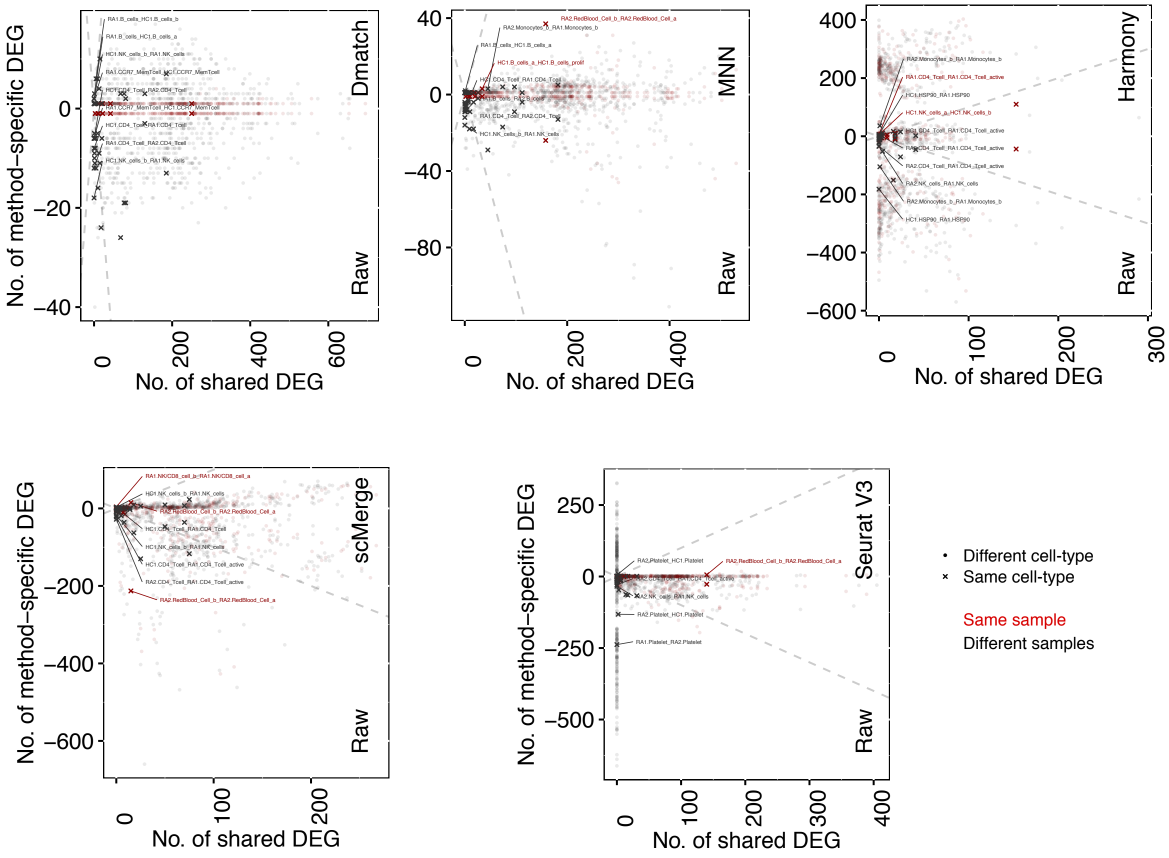


Supplementary Figure 5: Scatter plot showing the number of differentially expressed genes (DEG) that are identified using unaligned data (positive y-axis) or using data aligned using different methods (negative y-axis), versus the number of DEG that are identified in both datasets (x-axis). Each point represents a comparison between cell-type clusters from the same or different cell-type, or from the same or different sample. Successful removal of batch effect is supported by smaller numbers of DEGs resulting from aligned data in same cell-type comparisons, compared to unaligned data (smaller positive y-axis than negative y-axis for “x”’s). However, for most methods other than `Dmatch`, there were considerable variation in DEG between unaligned and aligned data for comparisons across clusters from the same sample (red “x”’s or points).

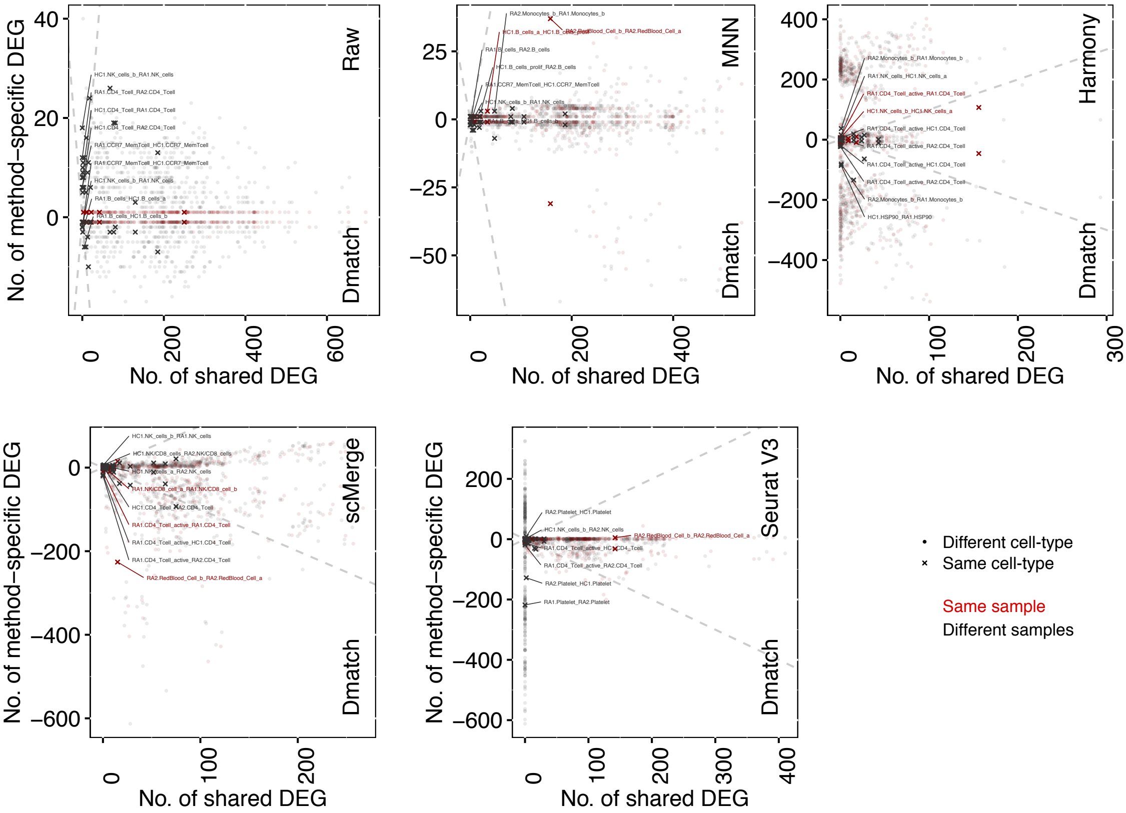


Supplementary Figure 6: Scatter plot showing the number of differentially expressed genes (DEG) that are identified using `Dmatch`-aligned data (negative y-axis) or using data aligned using different methods (positive y-axis), versus the number of DEG that are identified in both datasets (x-axis). Each point represents a comparison between cell-type clusters from the same or different cell-type, or from the same or different sample. We observed substantial differences between DEGs identified after alignment from `Dmatch` and other alignment methods.

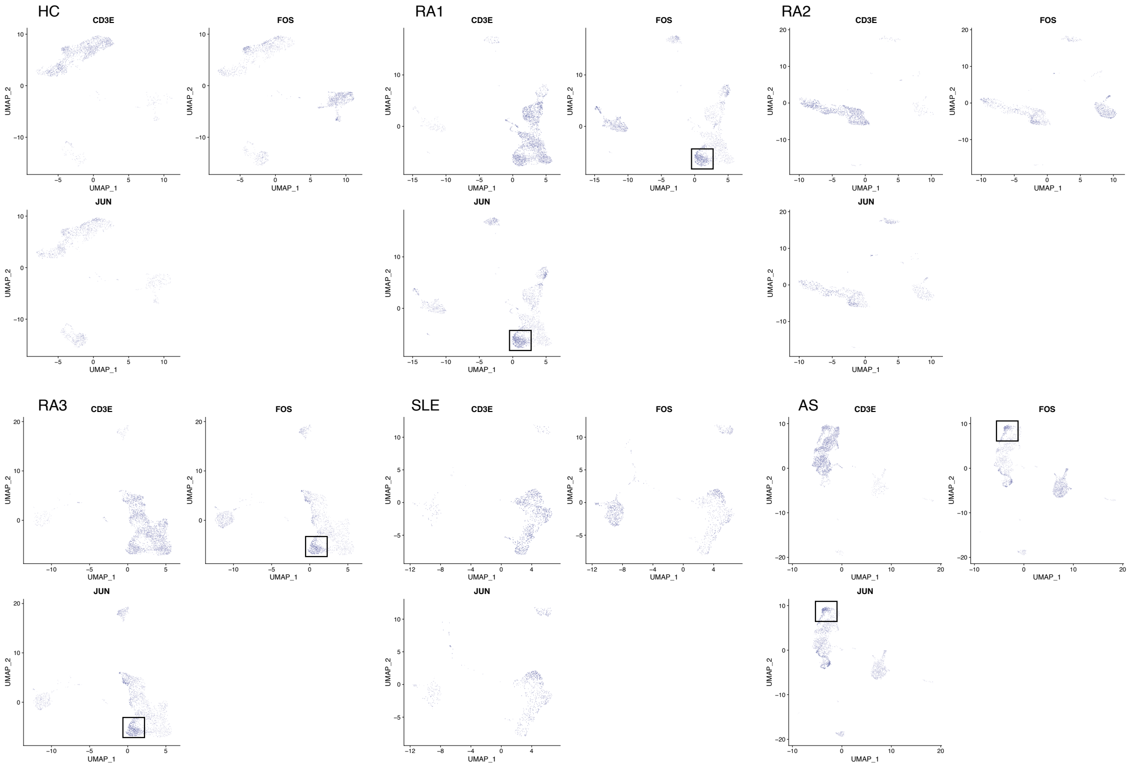


Supplementary Figure 7: UMAP of dimensionality reduction of PBMC scRNA-seq data from 6 PBMC samples, separately. We observed *JUN* and *FOS* in a subcluster of T-cells in PBMC of patients with autoimmune disease (RA1, RA3, AS) whose monocytes also expressed *IL1B*. Shades of purple represent marker expression in each cell relative to other cells in the same UMAP.

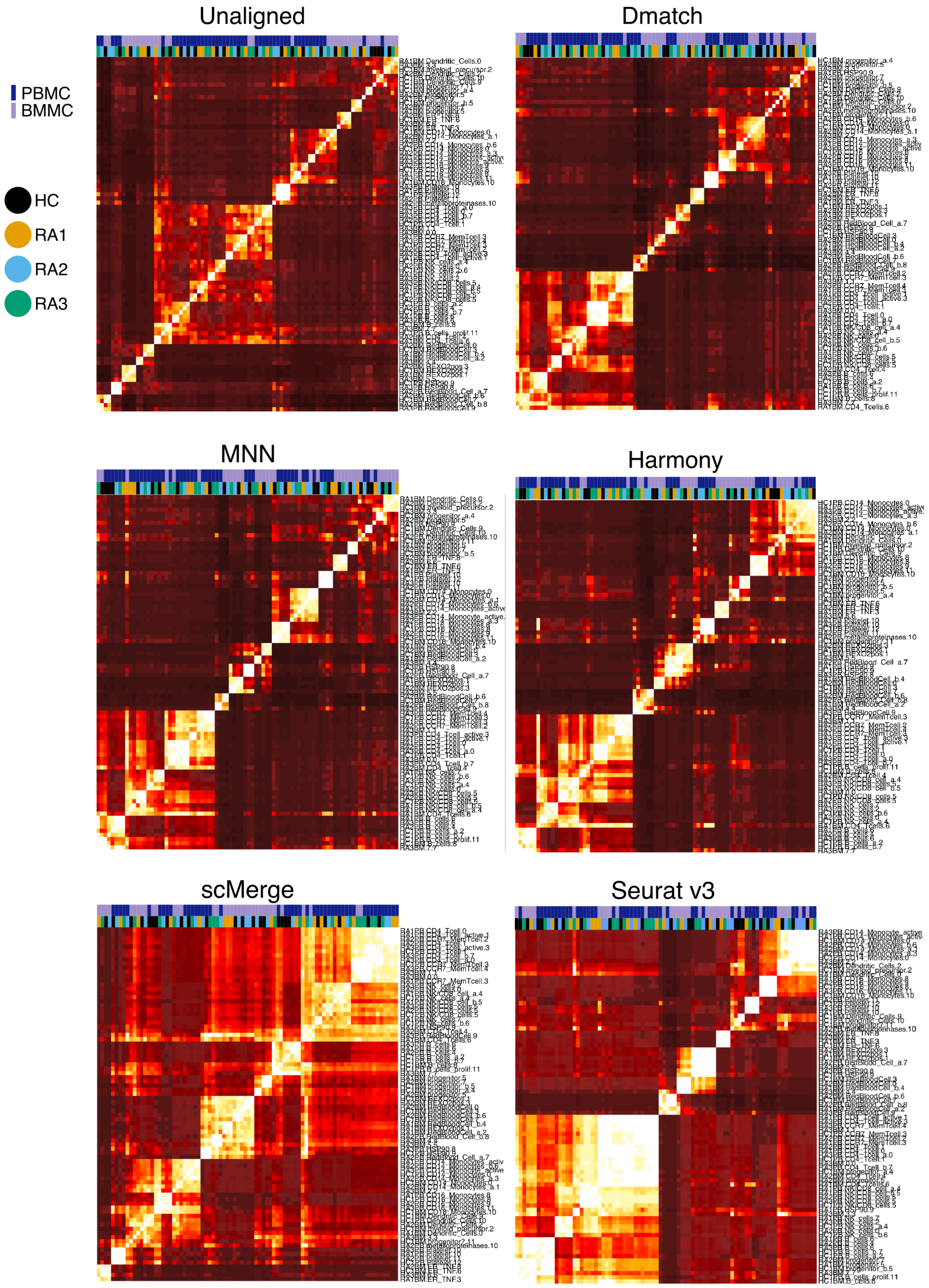


Supplementary Figure 8: Heatmaps showing the number of DEGs identified across Seurat clusters in PBMC and BMMC samples using aligned data from different methods.

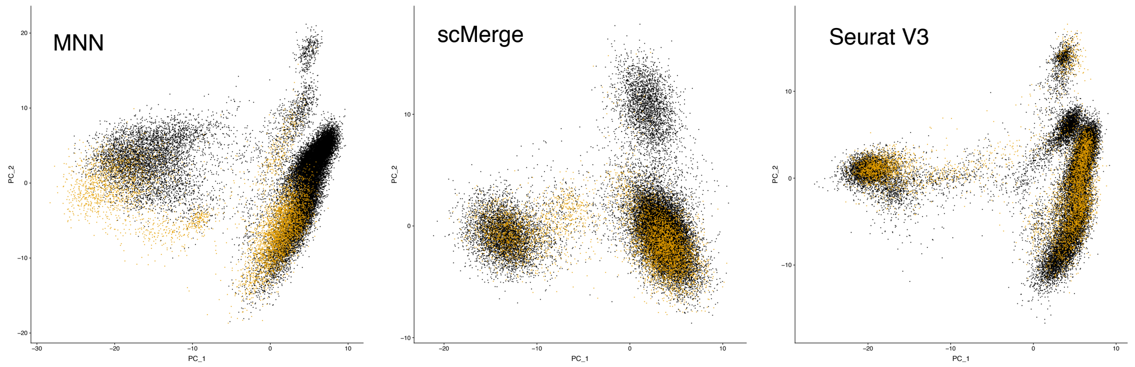


Supplementary Figure 9: PCA of scRNA-seq samples from PBMC and SF after alignment using different methods. Batch effect is reduced after alignment using all methods.

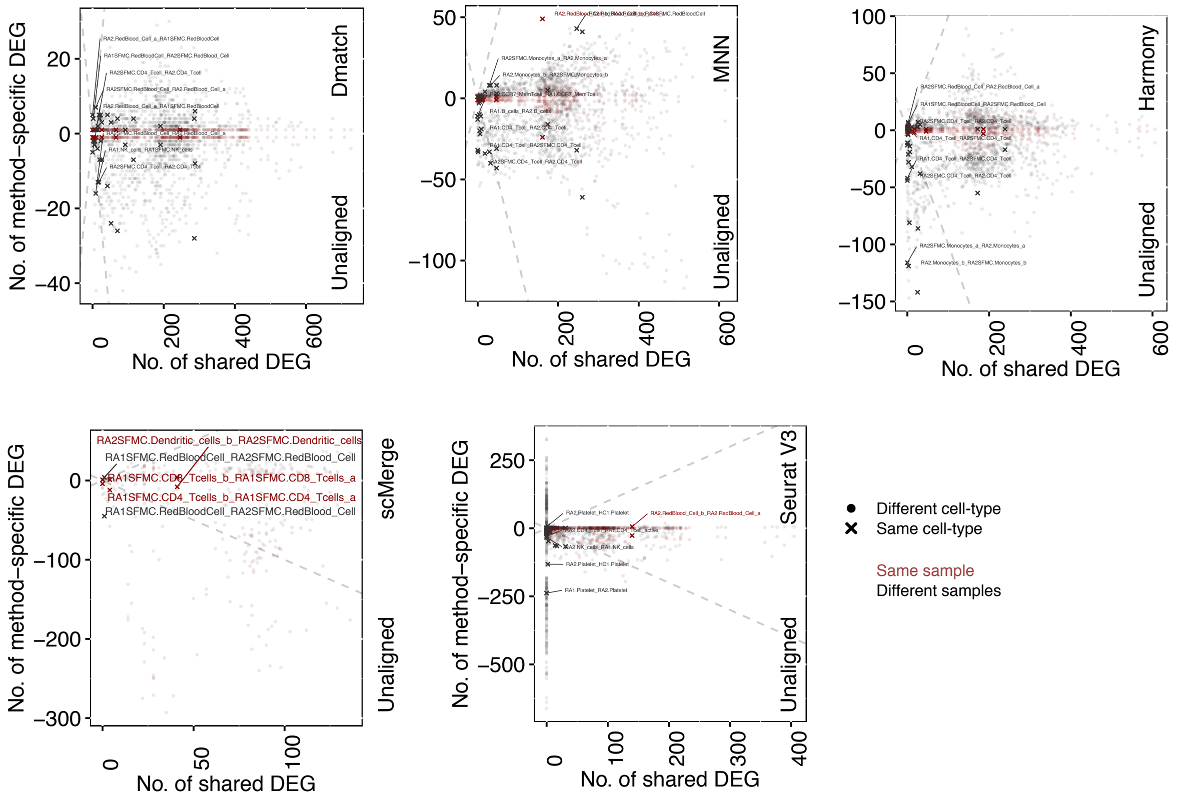


Supplementary Figure 10: Scatter plot showing the number of differentially expressed genes (DEG) that are identified using unaligned data (negative y-axis) or using data aligned using different methods (positive y-axis), versus the number of DEG that are identified in both datasets (x-axis). Each point represents a comparison between cell-type clusters from the same or different cell-type, or from the same or different sample. Successful removal of batch effect is supported by smaller numbers of DEGs resulting from aligned data in same cell-type comparisons, compared to unaligned data (smaller positive y-axis than negative y-axis for “x”’s).

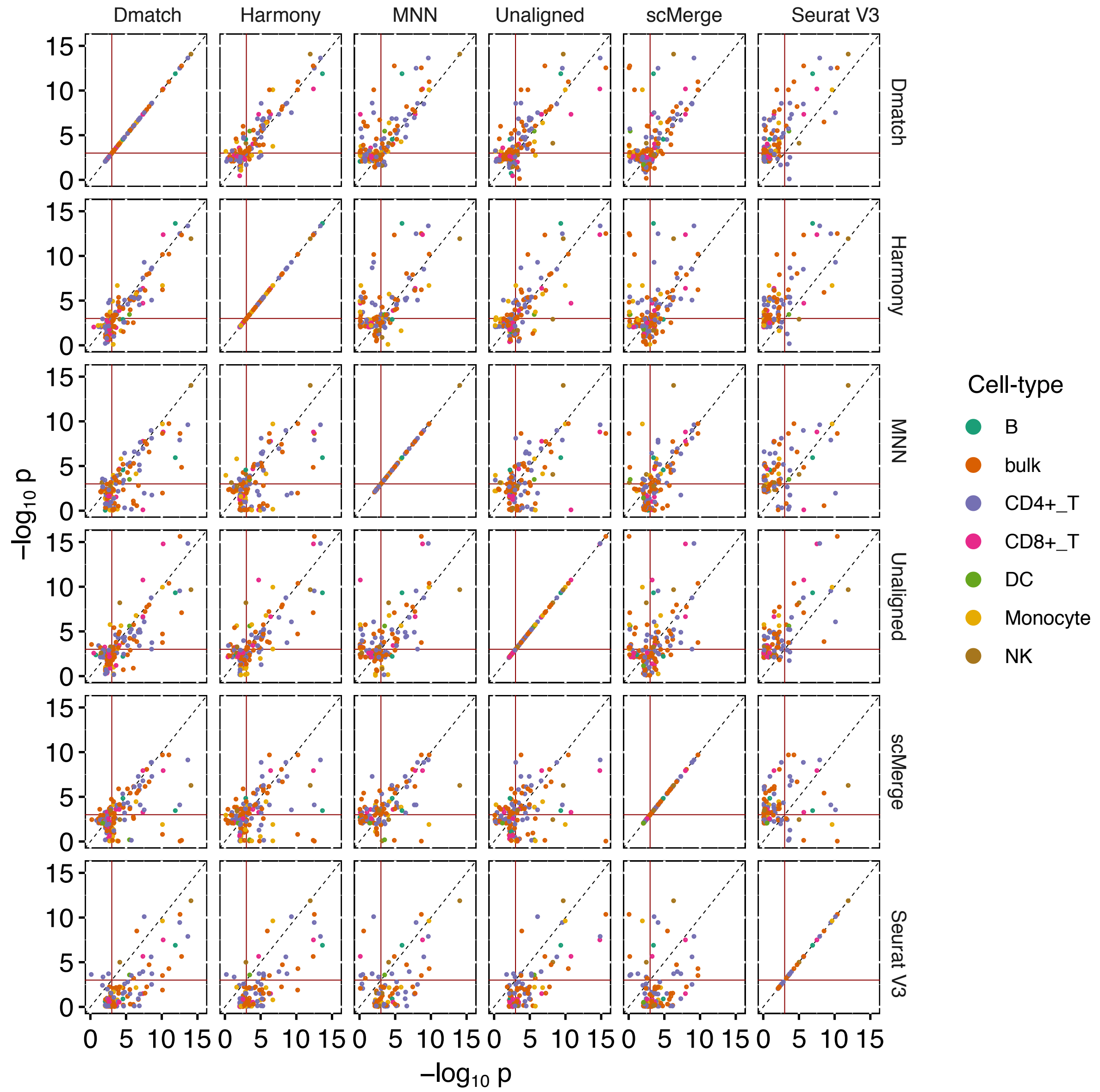


Supplementary Figure 11: Scatter plot showing the linear regression p-values comparing eQTLs (best gene-SNP pair) identified in different cell-types (colors) using unaligned or aligned data from different methods. While the p-values were generally highly correlated, in many cases they were more significant for `Dmatch`-aligned data compared to data aligned using other methods.

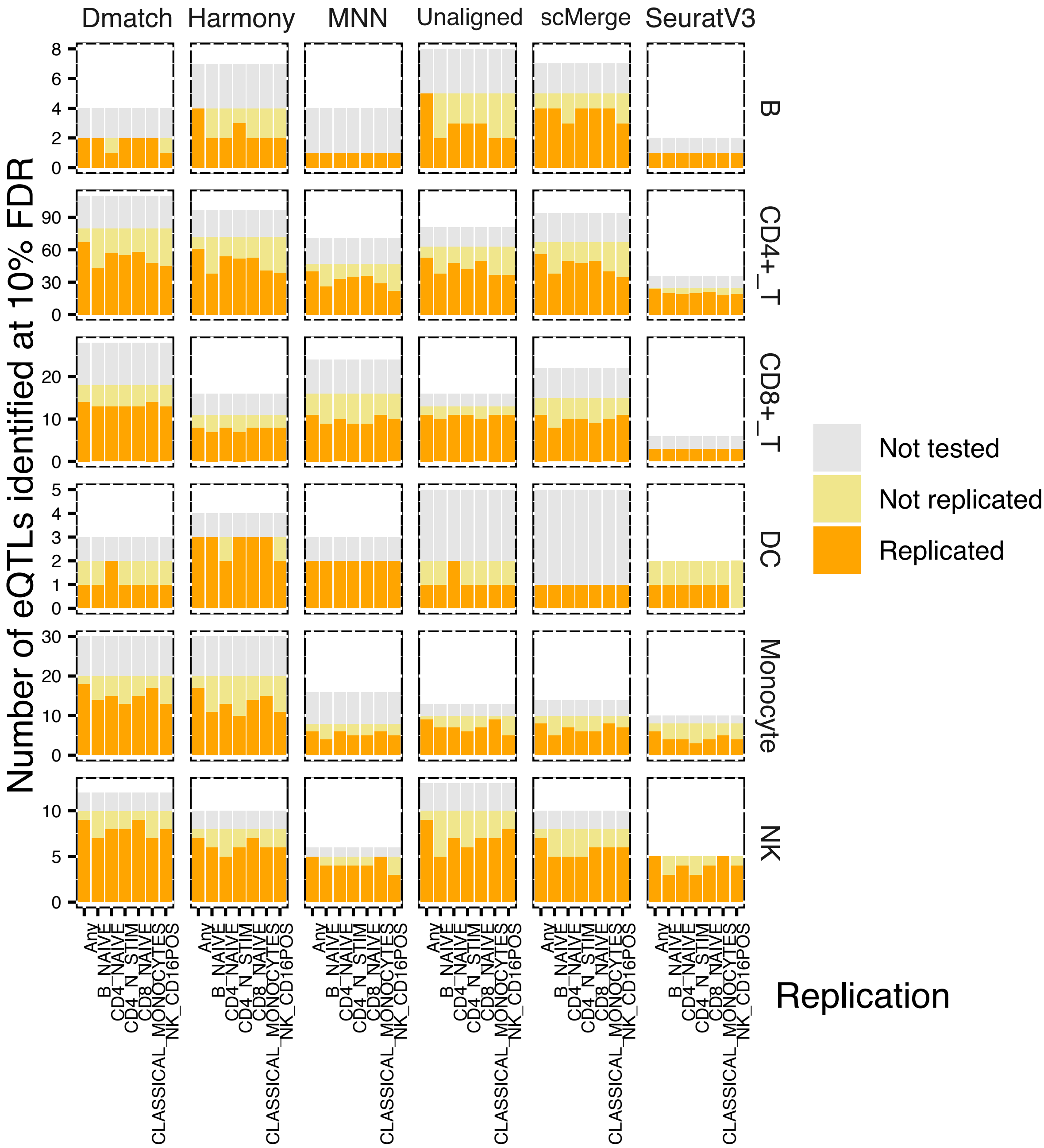


Supplementary Figure 12: Replication of eQTLs identified in a bulk RNA-seq study of immune cell-type eQTLs from the DICE consortium. The eQTLs that were not tested correspond to SNPs whose genotype could not be determined or to genes filtered out due to low expression.
