## Supplementary Table 1 for "Alignment of single-cell RNA-seq samples without over-correction using kernel density matching"

|  | HC | RA1 | RA2 | RA3 | AS | SLE |
| --- | --- | --- | --- | --- | --- | --- |
| Sex | Female | Female | Female | Female | Male | Female |
| Age (years) | 45 | 51 | 21 | 58 | 68 | 32 |
| Disease duration (years) | NA | 10 | 5 | 12 | 35 | 11 |
| ESR (mm/hour) | NA | 49 | 95 | 62 | 59 | 23 |
| CRP (mg/dL) | NA | 2.46 | 13.7 | 3.25 | 6.05 | Not detected |
| Clinical features | NA | RF+, Anti-CCP+, Anti-MCV+,  AKA+, ANA+, Anti-SSA+ | RF+, Anti-CCP+, Anti-MCV+,  AKA+ | RF+, Anti-CCP+, Anti-MCV+ | HLA-B27+ | ANA+, Anti-ds-DNA+, Anti-SSA+, Anti-Ro52+ Anti-nRNP/Sm+ |
| Complication | NA | None | Anemia | None | Hypertension | LN, LE |
| Treatment ^a^ | NA | None | None | None | None | Steroid |

ESR, erythrocyte sedimentation rate; CRP, C-reactive protein; RF, Rheumatoid factor; Anti-CCP, anti-cyclic citrullinated peptide antibody; Anti-MCV，anti-mutated citrullinated vimentin antibody; AKA, anti-keratin antibody; ANA, antinuclear antibody; Anti-ds-DNA, anti-double-stranded DNA antibody; Anti-SSA, anti-sjogren syndrome A antibody; Anti-Ro52, anti-Ro52 antibody; Anti-nRNP/Sm, anti ribonucleoprotein antibody; LN, lupus nephritis; LE, lupus encephalopathy. NA, not applicable.

^a^ Treatment within the last three months

**Supplementary Table 1.** Clinical characteristics of patients and healthy controls (HC)
