## Supplementary Table 2 for "Alignment of single-cell RNA-seq samples without over-correction using kernel density matching"

| Cell-type | No. of Phenotypic PC |
| --- | --- |
| B NAIVE | 12 |
| CD4 STIM | 10 |
| CD4 NAIVE | 10 |
| CD8 NAIVE | 6 |
| CLASSICAL MONOCYTES | 11 |
| NK CD16POS | 10 |

**Supplementary Table 2**: The number of phenotypic PCs used as covariates in our linear regression. The number of phenotypic PCs were chosen to maximize the number of eQTLs identified at 5% FDR.
